# Antimicrobial overproduction sustains intestinal inflammation by inhibiting *Enterococcus* colonization

**DOI:** 10.1101/2023.01.29.526128

**Authors:** Kyung Ku Jang, Thomas Heaney, Mariya London, Yi Ding, Frank Yeung, Defne Ercelen, Ying-Han Chen, Jordan Axelrad, Sakteesh Gurunathan, A. Marijke Keestra-Gounder, Matthew E. Griffin, Howard C. Hang, Ken Cadwell

## Abstract

Loss of antimicrobial proteins such as REG3 family members compromises the integrity of the intestinal barrier. Here, we demonstrate that overproduction of REG3 proteins can also be detrimental by reducing a protective species in the microbiota. Patients with inflammatory bowel disease (IBD) experiencing flares displayed heightened levels of secreted REG3 proteins that mediated depletion of *Enterococcus faecium* (*Efm*) from the gut microbiota. *Efm* inoculation of mice ameliorated intestinal inflammation through activation of the innate immune receptor NOD2, which was associated with the bacterial DL-endopeptidase SagA. Microbiota sensing by NOD2 in myeloid cells mediated IL-1β secretion and increased the proportion of IL-22-producing CD4^+^ T helper cells and innate lymphoid cells. Finally, *Efm* was unable to protect mice carrying a *NOD2* gene variant commonly found in IBD patients. Our findings demonstrate that inflammation self-perpetuates by causing aberrant antimicrobial activity that disrupts symbiotic relationships with gut microbes.

## INTRODUCTION

Secretion of antimicrobial proteins (AMPs) at barrier surfaces represents a ubiquitous and evolutionarily ancient defense strategy. AMPs such as defensins, cathelicidins, lysozyme, and C-type lectins are abundant in the mammalian gastrointestinal tract where they limit the overgrowth or invasion of microbes, typically by disrupting membrane integrity of their targets^1–3^. Although certain AMPs recognize structures associated with invasive microbes such as flagellin^4^, many display broad-spectrum activity, reflecting the diversity of pathogenic and non-pathogenic infectious agents in the gut. Regenerating family member 3 (REG3) C-type lectins REG3A and REG3G bind to the peptidoglycan cell wall of Gram-positive (+) bacteria and are essential for maintaining a balanced relationship with the gut microbiota^5, 6^. *Reg3γ* expression in mice is induced by toll-like receptor (TLR) signaling in the intestinal epithelium to achieve spatial segregation between the host and the microbiota^7–9^. REG3A (also known as HIP/PAP) is the main paralog that has been characterized in humans and may have a similar role as murine REG3G because transgenic expression of the human gene in mice alters microbiota composition^10^. In addition to inducing lysis of microbes, REG3 proteins contribute to intestinal homeostasis by regulating crypt regeneration and scavenging reactive oxygen species^10,11^

Dysregulated AMP production exacerbates intestinal injury in animal models and is associated with human inflammatory bowel disease (IBD)^12–17^, a chronic immune-mediated disorder of the gut involving complex interactions between genetic and environmental factors. IBD affects more than 6 million worldwide, and although immunomodulatory medications can offer substantial relief to patients, maintaining disease remission remains a major clinical challenge. Consistent with the link between AMPs and intestinal inflammation, an altered microbiota composition is a common feature of IBD^18–20^. Also, population genetic studies have implicated genes associated with host-microbe interactions^21, 22^. Coding region variants of the intracellular microbial sensor nucleotide-binding oligomerization domain-containing protein 2 (NOD2) are among the strongest genetic risk factors for IBD^23, 24^. In the presence of peptidoglycan byproducts, NOD2 mediates production of AMPs and cytokines downstream of NFκB activation^23^. Accordingly, *Nod2*-deficient mice are susceptible to enteric bacterial infections and display an altered microbiota composition depending on the animal facility^25–34^, and IBD patients harboring *NOD2* variants display impaired production of AMPs, especially defensins^35–37^. However, *NOD2* variants are found in only a subset of IBD patients, indicating that additional mechanisms may be involved. Also, IBD is associated with increased *REG3* expression^38, 39^, suggesting a different role for this family of AMPs in disease pathogenesis.

The *Enterococcus* genus includes Gram+ bacterial members of the gut microbiota that activate and are controlled by NOD2^40–42^, and are highly sensitive to killing by REG3A and REG3G^6, 9^. However, the relationship between enterococci and IBD is obfuscated by contradictory observations. *Enterococcus* species are opportunistic pathogens in hospital settings that cause life-threatening bloodstream infections^43^ and induce intestinal inflammation in *Il10* deficient mice^44, 45^. Yet, enterococci are consumed as probiotics for treatment of diarrheal diseases^46, 47^. Recent studies have provided a mechanistic basis for their probiotic properties by identifying a DL-endopeptidase, SagA, secreted by certain enterococci including *Enterococcus faecium* (*Efm*)^48–50^. Processing of peptidoglycan by SagA generates muropeptides, which potently activate NOD2 to induce an antimicrobial and inflammatory gene expression program that enhances colonization resistance towards enteric pathogens and antitumor immunity^41, 50, 51^. Interventions that promote antitumor immunity typically exacerbate intestinal inflammation^52^. Therefore, it is unclear whether NOD2 activation by enterococci would worsen or improve IBD.

In this study, we found IBD patients displayed uncontrolled production of REG3 proteins that deplete enterococci from the gut microbiota. Inoculation with *Efm* or a *Lactococcus* strain engineered to express *SagA* protected mice from intestinal injury. Protection was dependent on NOD2 activity in myeloid cells, which mediated an increase in lymphoid cells producing the regenerative cytokine IL-22. This beneficial effect of *Efm* colonization was abrogated in mice harboring the R702W variant of NOD2 associated with IBD. Our findings uncover a mechanism of perpetuating intestinal inflammation initiated by a harmful feedback loop involving excess AMP production and depletion of protective enterococci, which renders the host functionally NOD2-deficient.

## RESULTS

### REG3 proteins overproduced by IBD patients inhibit enterococci

Although *REG3A* mRNA was shown to be increased in inflamed intestinal tissues from IBD patients^38, 39^, only a small number of samples were analyzed in these prior studies, and it is unclear whether gene expression is associated with REG3 protein secretion and activity. Thus, the functional consequence has not been investigated. To measure antimicrobial activity in the gut, we collected stool from 56 IBD patients experiencing disease flares (Supplemental table 1). An equal number of non-IBD patients (NIBD) experiencing gastrointestinal symptoms, such as diarrhea, were used as a control cohort (Supplemental table 2). Stool extracts were prepared by 2-step filtrations (see METHOD DETAILS) and subjected to Western blot analysis of the two human REG3 family members, REG3A and REG3G (Figure 1A). We also examined two additional paralogs: REG1A, which is a pancreatic growth factor and should not be co-regulated with REG3 proteins^53^, and REG4, a potential AMP that is less characterized^54, 55^. As we expected, pancreatic growth factor REG1A is equally present in NIBD and IBD (Figure 1B and C). In contrast, REG3A was detected in a higher proportion of IBD patients compared with NIBD patients, and REG3G and REG4 were exclusively present in IBD samples (Figure 1B and C). An ELISA assay confirmed higher REG3A concentration in stool extracts from IBD compared with those of NIBD (Figure 1D). These results indicate that the presence of REG3 proteins is more prevalent and enriched in IBD patients.

**Figure 1.**
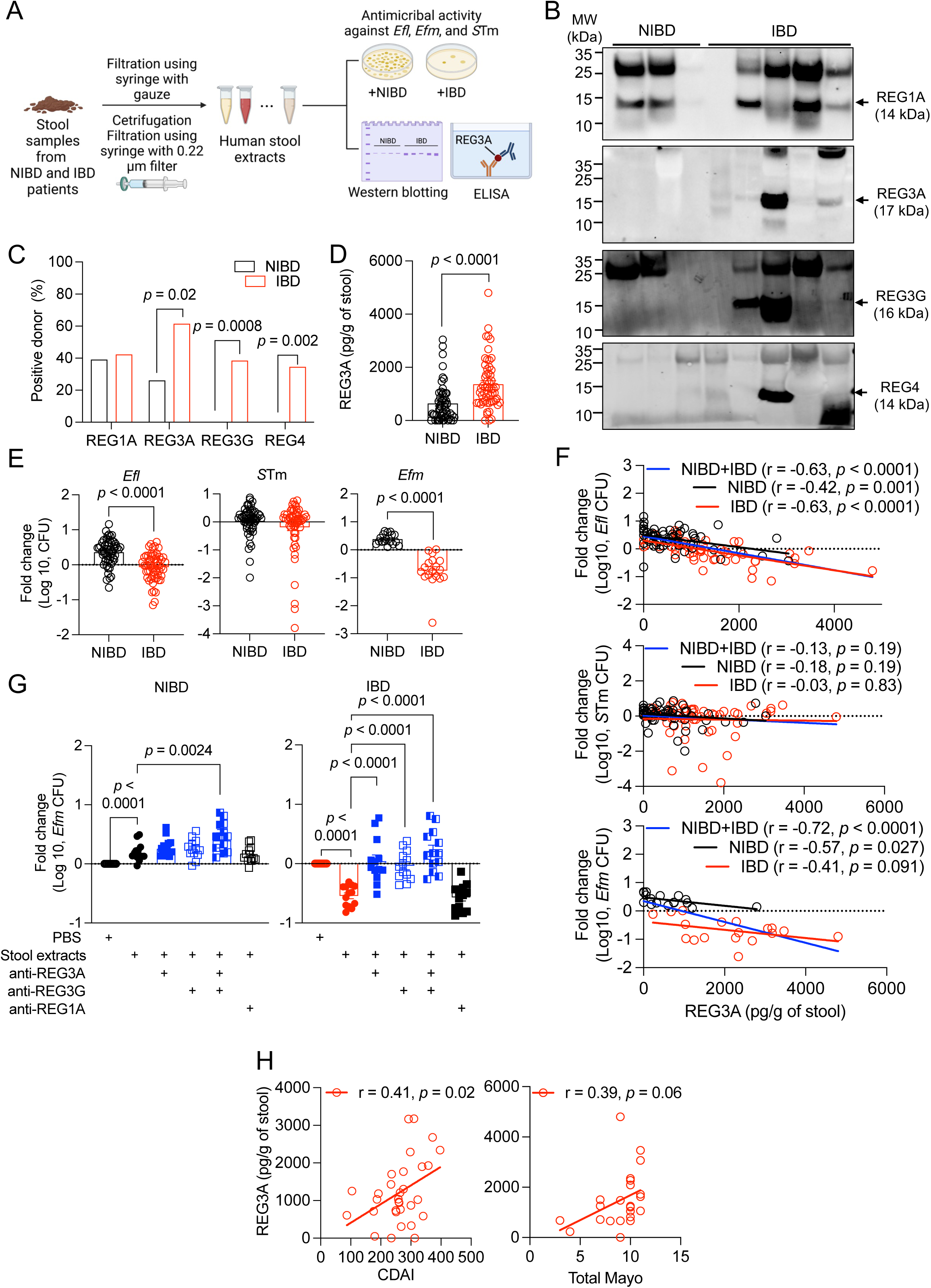
Increased REG3A and REG3G in IBD patient stool inhibit *Enterococcus*. **A)** Schematic of stool extract preparation and analysis. **B)** Western blots of REG1A, REG3A, REG3G, and REG4 in stool extracts from representative 3 non-IBD (NIBD) and 5 IBD patients. **C)** Proportion of stool specimens from individual NIBD (n = 23) and IBD (n = 26) patients in which REG1A, REG3A, REG3G, or REG4 were detectable by western blot. **D)** Quantification of REG3A in NIBD and IBD patients-derived stool extracts by ELISA. **E)** Fold changes in colony forming units (CFU) of *Enterococcus faecalis* (*Efl*) and *Salmonella* Typhimurium (*S*Tm) following 24 hrs of culture in stool extracts from NIBD (n = 56) and IBD (n = 56) patients. Results obtained with *Efl* were confirmed with *E. faecium* (*Efm*) (n = 18 for NIBD and n = 19 for IBD). **F)** Correlation between REG3A concentration and CFU fold changes of *Efl* (upper), *S*Tm (middle), and *Efm* (lower) cultured in stool extract from NIBD and IBD patients. **G)** Fold change in *Efm* CFU at 24 hr-post culture in NIBD (left) and IBD (right) patient-derived stool extract or PBS in the presence of anti-REG1A, REG3A, and REG3G antibodies. N = 13 for NIBD and IBD. **H)** Correlation between REG3A concentration and Crohn’s disease activity index (CDAI) for CD patients (n = 31, left) or total Mayo score for ulcerative colitis (UC) patients (n = 24, right). Data points in D-H represent individual patients. Bars represent mean ± SEM and at least three independent experiments were performed. NIBD, non-IBD; *Efl*, *E. faecalis*; *S*Tm, *S.* Typhimurium; *Efm*, *E. faecium*; CDAI, Crohn’s disease activity index; UC, ulcerative colitis; r, Pearson correlation coefficient. Indicated *p* values by Fisher’s exact test in C, unpaired *t* test, two-tailed in D, E, and G, and simple linear regression analysis in F and H.

Next, we examined the antimicrobial activity of the IBD and NIBD-derived stool extracts against *Enterococcus faecalis* (*Efl*) and *Salmonella* Typhimurium (*S*Tm) as representative Gram+ and – enteric bacteria, respectively. The number of *Efl* colonies recovered after culturing in IBD stool extracts was significantly lower compared with NIBD stool extracts, whereas *S*Tm numbers were similar in both groups. (Figure 1E). We confirmed this antimicrobial activity of IBD stool extracts with another *Enterococcus* species, *Efm* (Figure 1E). Consistent with Gram+ bacterial targeting, REG3A concentration negatively correlated with *Efl* and *Efm* colony forming units (CFUs) following treatment with NIBD and IBD stool extracts, but not with *S*Tm CFUs (Figure 1F). To determine whether the antimicrobial activity in stool samples was dependent on REG3 proteins, we added anti-REG3A or -REG3G antibodies, individually or together, to *Efm* cultured in the stool extracts. Less *Efm* was recovered when grown in IBD stool extracts compared with PBS control, and this antimicrobial activity was inhibited by antibodies against REG3A or REG3G, but not REG1A control antibodies (Figure 1G). Unlike IBD samples, an increased number of *Efm* was recovered when cultured in NIBD stool extracts compared with PBS control, and adding anti-REG3A and REG3G antibodies together (but not individually) enhanced growth, suggesting low levels of antimicrobial activity was present in these samples (Figure 1G). Thus, *Efm* growth impairment in human stool extracts is attributed to REG3A and REG3G.

We next segregated the data based on sex, disease type and severity, and age. IBD stool extracts from both males and females displayed higher amounts of REG3A compared with NIBD samples (Supplemental figure 1A). IBD consists of two major forms, Crohn’s disease (CD) and ulcerative colitis (UC); REG3A concentrations were similar in stool from CD and UC patients (Supplemental figure 1B). REG3A concentration did not differ among age groups in NIBD patients, however, displayed a modest but significant inverse relationship with age among IBD patients (Supplemental figure 1C). When controlling for age, we found that IBD patients younger than age 60 displayed increased levels of REG3A compared with matched NIBD controls, and this difference was lost in the older age group (Supplemental figure 1D). Disease severity measurements of CD (Crohn’s disease activity index, CDAI) and UC (total Mayo) displayed a positive correlation with REG3A concentrations (Figure 1H). Thus, the combined results suggest that the increase in REG3 in stool from IBD patients, proportional to disease severity, inhibits *Enterococcus* species.

### *Enterococcus* and *Efm* are lost from the gut microbiota in IBD patients

The above results raise the possibility that *Enterococcus* is depleted in the gut microbiota of IBD patients. Analysis of stool by 16S rRNA sequencing indicated that the overall microbiota composition of NIBD and IBD were indistinguishable according to multiple metrics of alpha and beta diversities (Figure 2A and B). These results further confirm that our NIBD cohort is an appropriate control for IBD patients. However, >80% of the sequencing reads aligned to *Enterococcus* in one of the NIBD samples, indicating that this individual had an overabundance of *Enterococcus* and thus, we excluded this patient from subsequent analyses. Consistent with our *in vitro* findings (Figure 1), the relative abundance of *Enterococcus* in IBD patients was lower than that in NIBD patients (Figure 2C).

**Figure 2.**
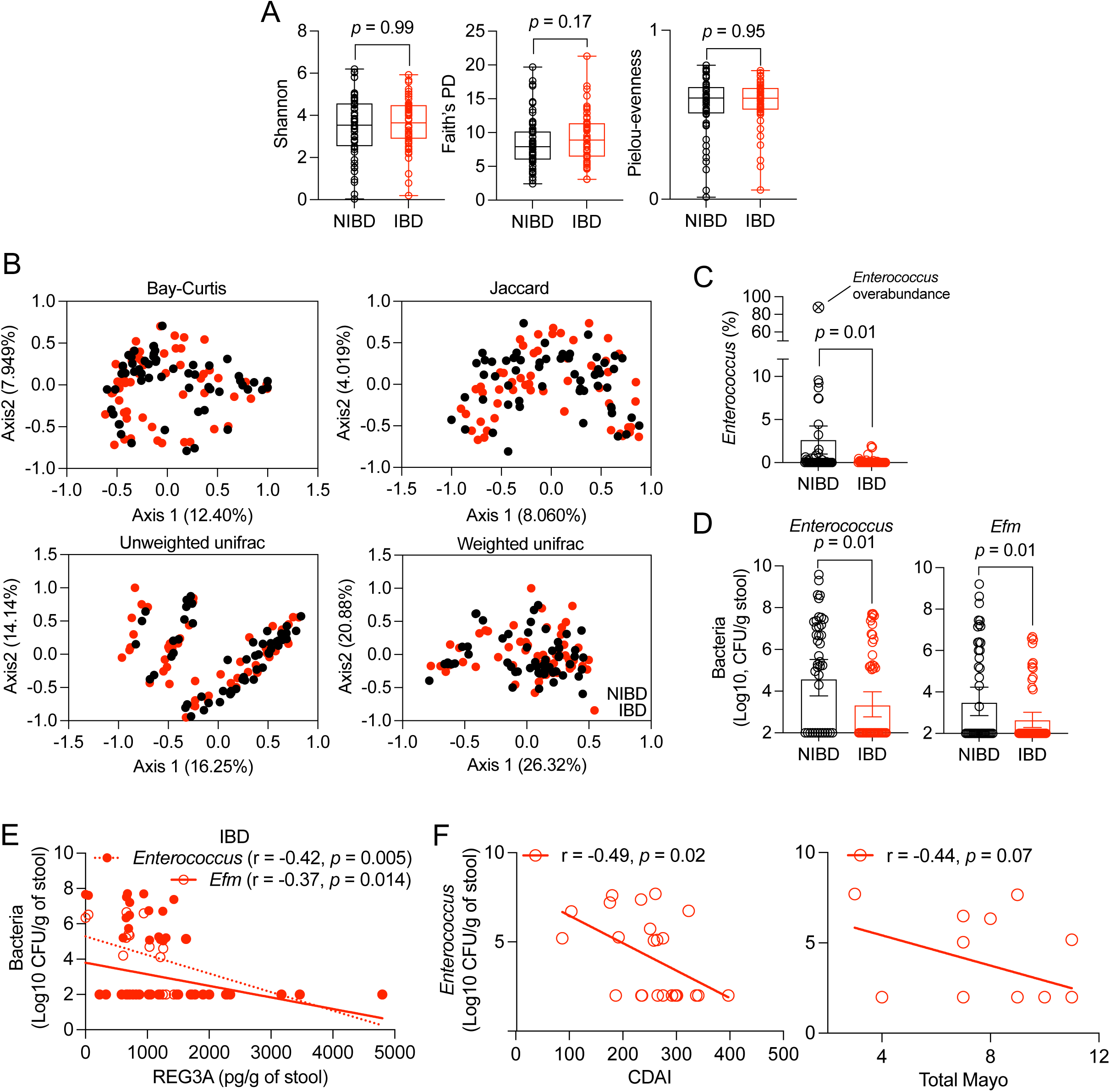
*Enterococcus* and *Efm* are depleted in gut microbiota from IBD patients. **A and B)** 16S rRNA sequencing of stool from NIBD and IBD patients from Figure 1. Alpha diversity values calculated as Shannon (right), Faith’s phylogenetic diversity (PD) (middle), and Pielou-evenness (right) indices (A). Principle coordinate analyses of beta diversity determined by Bay-Curtis, Jaccard, and Unweighted and Weighted unifrac methods (B). **C)** Proportion of sequencing reads representing *Enterococcus* in NIBD and IBD patient stool. One NIBD sample contained >80% *Enterococcus* indicative of an overabundance, which is shown as a reference on the graph but excluded from statistical analysis and downstream assays. **D)** Total *Enterococcus* (left) and *Efm* (right) CFUs in stool from NIBD (n = 39) and IBD (n = 43) patients. **E)** Correlation between REG3A concentration and the burden of total *Enterococcus* and *Efm* in IBD stool. **F)** Correlation between the burden of total *Enterococcus* and CDAI of CD patients (n = 24, left) and total Mayo score of UC patients (n = 18, right). Data points represent individual patients. Bars represent mean ± SEM and at least three independent experiments were performed. r, Pearson correlation coefficient. Indicated *p* values by Kruskal-Wallis test in A, unpaired *t* test, two-tailed in C and D, and simple linear regression analysis in E and F.

To confirm our 16S sequencing result, we plated NIBD and IBD stool on media that allows enumeration of enterococci or that is selective for *Efm*. The burden and detection rate of total *Enterococcus* and *Efm* in IBD stools were lower than in NIBD stools (Figure 2D and Supplemental figure 1E). *Efm* accounted for less than 20% of the total enterococci present in IBD stool compared with 49% in NIBD stool, indicating that *Efm* was particularly vulnerable to depletion in IBD patients. (Supplemental figure 1F). REG3A concentration displayed a negative correlation with total *Enterococcus* and *Efm* burden in IBD patient-derived stool (Figure 2E) as well as with CDAI and total Mayo score (Figure 2F). These results, combined with *in vitro* bacterial culturing data, indicate that levels of enterococci including *Efm* are reduced by the increased REG3 proteins in IBD patients.

### Enterococci protect against intestinal injury in mice

We next wanted to examine the relationship between *Enterococcus* dynamics and intestinal inflammation. While performing experiments to determine the appropriate conditions for administering dextran sulfate sodium (DSS) in drinking water, a widely used model of intestinal chemical injury, we observed extreme differences in susceptibility between wild-type (WT) C57BL/6J (B6) mice bred in two rooms within the same institutional vivarium. Mice bred in room 6 receiving 5% DSS displayed higher lethality, weight loss, and disease score compared with mice raised in room 13 (Figure 3A-C). Lipocalin-2 (LCN2), a marker of inflammation^56^, was also more abundant in stool from DSS-treated room 6 mice compared with room 13 mice (Figure 3D). Susceptibility to DSS can diverge between B6 mice, even those bred in the same animal facility, due to differences in microbiota composition that arise from separate parental lineages^57, 58^. Because we were performing these experiments with the goal of investigating enterococci, we plated stool specimens to examine their presence in the microbiota and observed a 2-log higher burden of endogenous enterococci in room 13 mice than room 6 mice (Figure 3E). Similar to the inverse relationship with inflammation observed in human samples, *Enterococcus* levels decreased following DSS-induced intestinal inflammation in mice (Figure 3F). In all subsequent experiments, we used mice from room 6.

**Figure 3.**
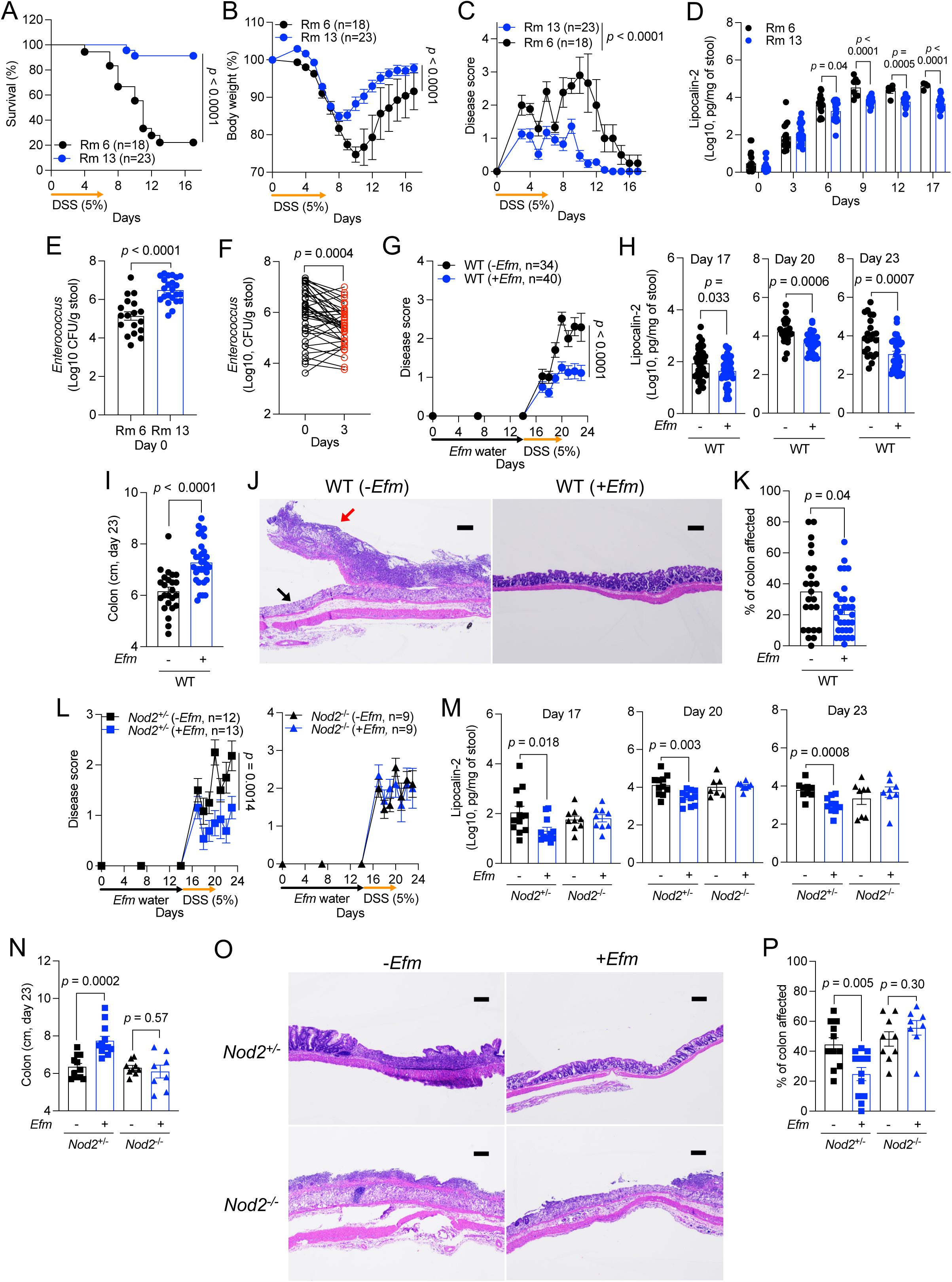
*Enterococcus* protects against intestinal injury in a NOD2-dependent manner. **A-F)** Dextran sulfate sodium (DSS)-induced intestinal injury of wild-type (WT) B6 mice bred in room 6 or 13 in the NYU animal facility. The mice were examined for survival (A), changes in body weight (B) and disease score (C), and lipocalin-2 (LCN2) in stool samples at the indicated time points (D). Endogenous *Enterococcus* burden was measured on day 0 (E) and changes in the *Enterococcus* burden on day 3 (F). **G-I)** DSS-treatment of WT mice from room 6 following administration of *Efm* in drinking water or control. The mice were examined for changes in disease score (G), LCN2 in stool at the indicated time points (H), and colon length (I) on day 23. **J and K)** Representative images of hematoxylin and eosin (H&E)-stained sections of the colon (J) and quantification of the proportion of colon affected (K). In the left panel of J, black arrow indicates destroyed colonic epithelium replaced with a diffuse ulcer and pus, and red arrow indicates acute inflammation with primary neutrophils, lymphocytes, plasma cells, and debris of dead cells. A breakdown of histological signs of inflammation by individual mice is listed in Supplemental table 3 for this and other figures. **L-N)** DSS-treatment of *Nod2^+/-^* and *Nod2^-/-^* mice from room 6 following administration of *Efm* in drinking water or control were examined for changes in disease score (L), LCN2 in stool at the indicated time points (M), and colon length on day 23 (N). **O and P)** Representative images of H&E-stained sections of the colon (O) and quantification of the proportion of colon affected (P). Bars, 200 μm. Data points in D-F, H, I, K, M, N, and P represent individual mice. Data points in B, C, G, and L represent mean ± SEM. Bars represent mean ± SEM and at least three independent experiments were performed. DSS, dextran sulfate sodium; Rm 6, room 6; Rm 13, room 13; WT, wild type. Indicated *p* values by log-rank Mantel-Cox test in A, two-way ANOVA test in B, C, G, and L, unpaired *t* test, two-tailed in D, E, H, I, K, M, N, and P, and paired *t* test, two-tailed in F.

To determine whether *Enterococcus* colonization protects against intestinal injury, we inoculated mice by administering *Efm* in the drinking water (10^9^ CFU/ml) for 2 weeks^59^ and then switched to water containing 5% DSS for an additional 6 days. Body weight and fecal LCN2 concentration were unchanged in mice inoculated with *Efm* in the absence of DSS treatment (Supplemental figure 2A and B), confirming that *Efm* colonization does not induce inflammation. In mice receiving DSS, both total *Enterococcus* and *Efm* levels decreased during the treatment and partially recovered after cessation (Supplemental figure 2C and D). *Efm*-colonized WT B6 mice treated with DSS displayed less body weight reduction, disease score, and fecal LCN2 compared with DSS-treated mice that were not inoculated with *Efm* (Figure 3G and H and Supplemental figure 2E and F). Mice were sacrificed on day 23 to determine whether improvement of these disease features reflected reduced intestinal inflammation. *Efm-*colonized mice displayed less shortening of the colon (a marker of intestinal inflammation) and histopathologic changes compared with uncolonized WT mice (Figure 3I-K and Supplemental table 3). Inoculating mice with *Efm* after DSS treatment also mitigated these signs of disease (Supplemental figure 3A-G and Supplemental table 3). Therefore, *Efm* colonization confers protection against DSS-induced inflammation.

Given that *Efm* protects against enteric pathogens in a NOD2-dependent manner^41^, we asked whether *Efm* would lose its beneficial properties when administered to *Nod2*-deficient mice during intestinal injury. The colonization patterns of *Efm* in *Nod2^+/-^* and *Nod2^-/-^* mice were consistent with those in WT mice (Supplemental figure 2G). In contrast to *Nod2^+/-^* mice in which we reproduced the protective effect of the bacterium, *Efm* administration to *Nod2^-/-^* mice did not improve weight loss, disease score, fecal LCN2 concentration, colon shortening, and intestinal histopathology (Figure 3L-P, Supplemental figure 2H and I, and Supplemental table 3). These findings indicate that the anti-inflammatory effect of *Efm* is dependent on *Nod2*.

### SagA mediates NOD2-dependent protection against intestinal injury

*Efm* encodes a secreted DL-endopeptidase SagA that processes peptidoglycan to generate NOD2-stimulating muropeptides^41, 50^. Because SagA is essential for growth of *Efm*^60^, we tested the role of SagA by orally inoculating mice with *Lactococcus lactis* (*Lls*) strains expressing wild-type SagA (*Lls*-*SagA^WT^*) or catalytically inactive (C443A) SagA (*Lls-SagA^C^*^443^*^A^*), then treating with 3% DSS (Supplemental figure 4A). A lower concentration of DSS was used because mice were pre-treated with antibiotics (abx) to facilitate *Lls* colonization, and abx increases susceptibility to DSS ^61, 62^. We included conditions in which mice were inoculated with PBS or the parental unmodified *Lls* strain to control for effects of abx treatment and *Lls* colonization, respectively. *Lls* strains displayed a similar degree of colonization (Supplemental figure 4B and D). Mice given *Lls*-*SagA^WT^* displayed enhanced survival compared with all the other conditions, and other disease parameters were generally improved (Figure 4A-D and Supplemental figure 4C). *Lls*-*SagA^WT^* ameliorated disease in *Nod2^+/^*^-^ mice but not *Nod2^-/-^* mice (Figure 4E-H and Supplemental figure 4E). If protection is mediated through peptidoglycan processing by secreted SagA, bacterial colonization would potentially be dispensable. To test this possibility, we administered filtered culture supernatants from *Lls* strains to mice without abx treatment for a week and then switched to water containing 5% DSS (Supplemental figure 4F). In contrast to supernatant collected from the parental *Lls* and *Lls-SagA^C^*^443^*^A^* cultures, or sterile broth, supernatant from *Lls*-*SagA^WT^* was sufficient for increasing survival, lessening body weight reduction, and decreasing disease score and fecal LCN2 concentration of DSS-treated mice (Figure 4I and J and Supplemental figure 4G and H). These protective effects of *Lls*-*SagA^WT^* culture supernatant were all NOD2-dependent (Figure 4K and L and Supplemental figure 4I and J). Taken together, our findings indicate that activating NOD2 downstream of SagA protects against intestinal injury.

**Figure 4.**
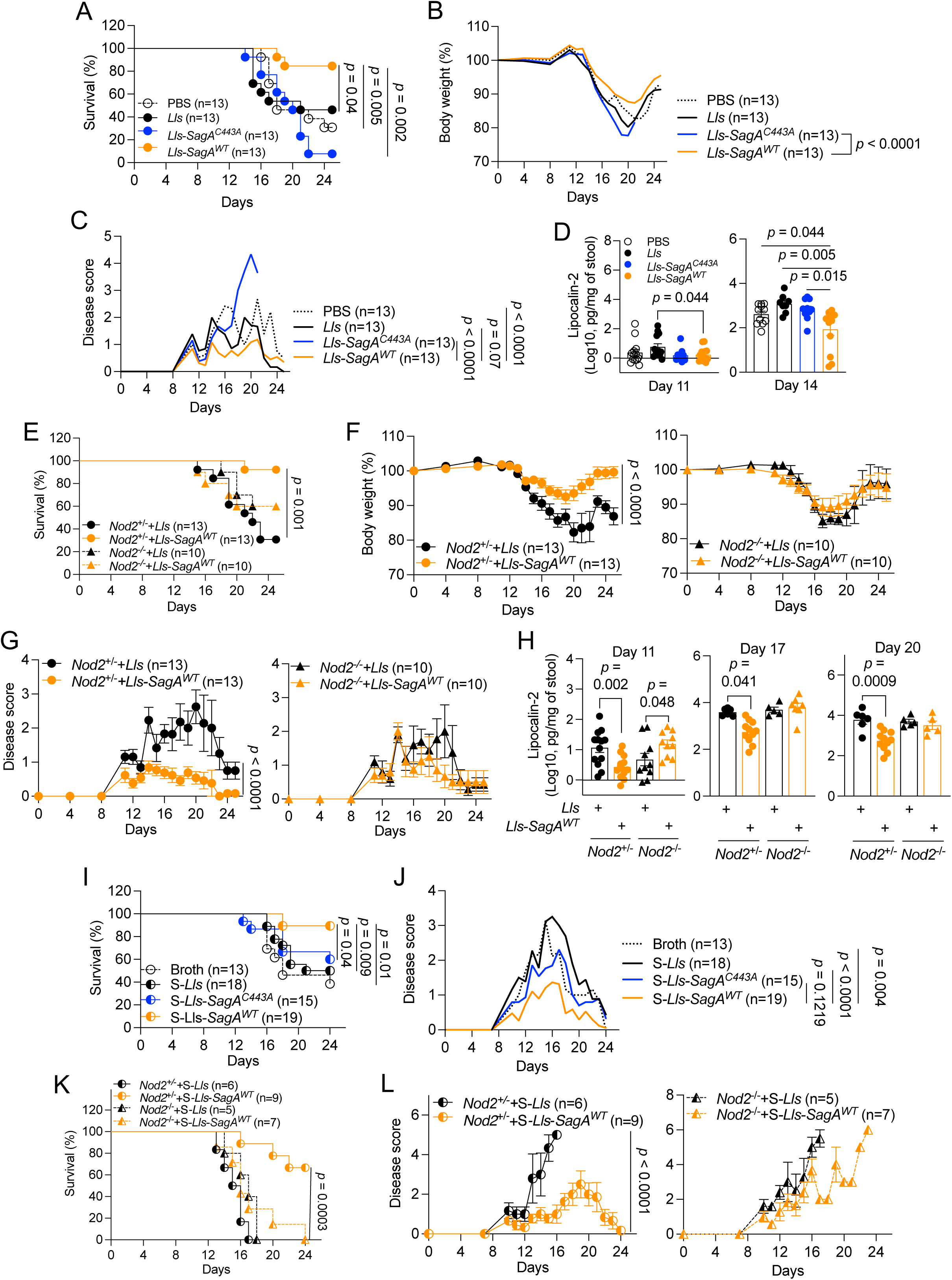
SagA mediates NOD2-dependent protection against intestinal injury. **A-D)** Antibiotics (ampicillin + streptomycin, abx)-treated WT mice orally inoculated with PBS, *Lactococcus lactis* (*Lls*), *Lls*-*SagA^WT^*, or *Lls*-*SagA^C443A^* and then given DSS were examined for survival (A), changes in body weight (B) and disease score (C), and LCN2 presence in stool samples at the indicated time points (D). **E-H)** Abx-treated *Nod2*^+/-^ and *Nod2*^-/-^ mice inoculated with *Lls* or *Lls*-*SagA^WT^* and then given DSS were examined for survival (E), changes in body weight (F) and disease score (G), and fecal LCN2 at the indicated time points (H). **I and J)** DSS-treated WT mice receiving broth or culture supernatants of *Lls* (S-*Lls*), *Lls*-*SagA^WT^* (S-*Lls*-*SagA^WT^*), or *Lls*-*SagA^C443A^* (S-*Lls*-*SagA^C443A^*) as drinking water were examined for survival (I) and changes in body weight (J). **K and L)** DSS-treated *Nod2*^+/-^ and *Nod2*^-/-^ mice receiving S-*Lls* or S-*Lls*-*SagA^WT^* as drinking water were examined for survival (K) and changes in disease score (L). All mice were bred in room 6. Data points in D and H represent individual mice. Lines in B, C, and J represent mean. Data points in F, G, and L represent mean ± SEM. Bars represent mean ± SEM and at least two independent experiments were performed. *Lls*, *L. lactis*; S, supernatant. Indicated *p* values by log-rank Mantel-Cox test in A, E, I, and K, two-way ANOVA test in B, C, F, G, J, and L, and unpaired *t* test, two-tailed in D and H.

### NOD2 in myeloid cells is indispensable for *Efm*-mediated protection

Myeloid and intestinal epithelial cells are among the cell types in which NOD2 function has been most extensively investigated^63–65^. Therefore, we developed *Nod2^fl/fl^;LysM-Cre* and *Villin-Cre* mice that are *Nod2*-deficient in myeloid and intestinal epithelial cells, respectively, and confirmed the presence of *Efm* in stool samples following inoculation in all groups (Supplemental figure 5A and C). Unlike *Nod2^fl/fl^* controls in which signs of disease were ameliorated, *Efm* colonization of *Nod2^fl/fl^;LysM-Cre^+^* mice did not improve lethality, body weight reduction, disease score, and LCN2 present in stool samples (Figure 5A-D and Supplemental figure 5B). In contrast, *Efm* colonization reduced mortality, body weight reduction, disease score, and fecal LCN2 concentration in *Nod2^fl/fl^;Villin-Cre^+^* mice to a similar extent as *Nod2^fl/fl^* controls (Figure 5E-H and Supplemental figure 5D). These findings indicate that NOD2 is required in myeloid cells and dispensable in the intestinal epithelium for *Efm*-mediated protection.

**Figure 5.**
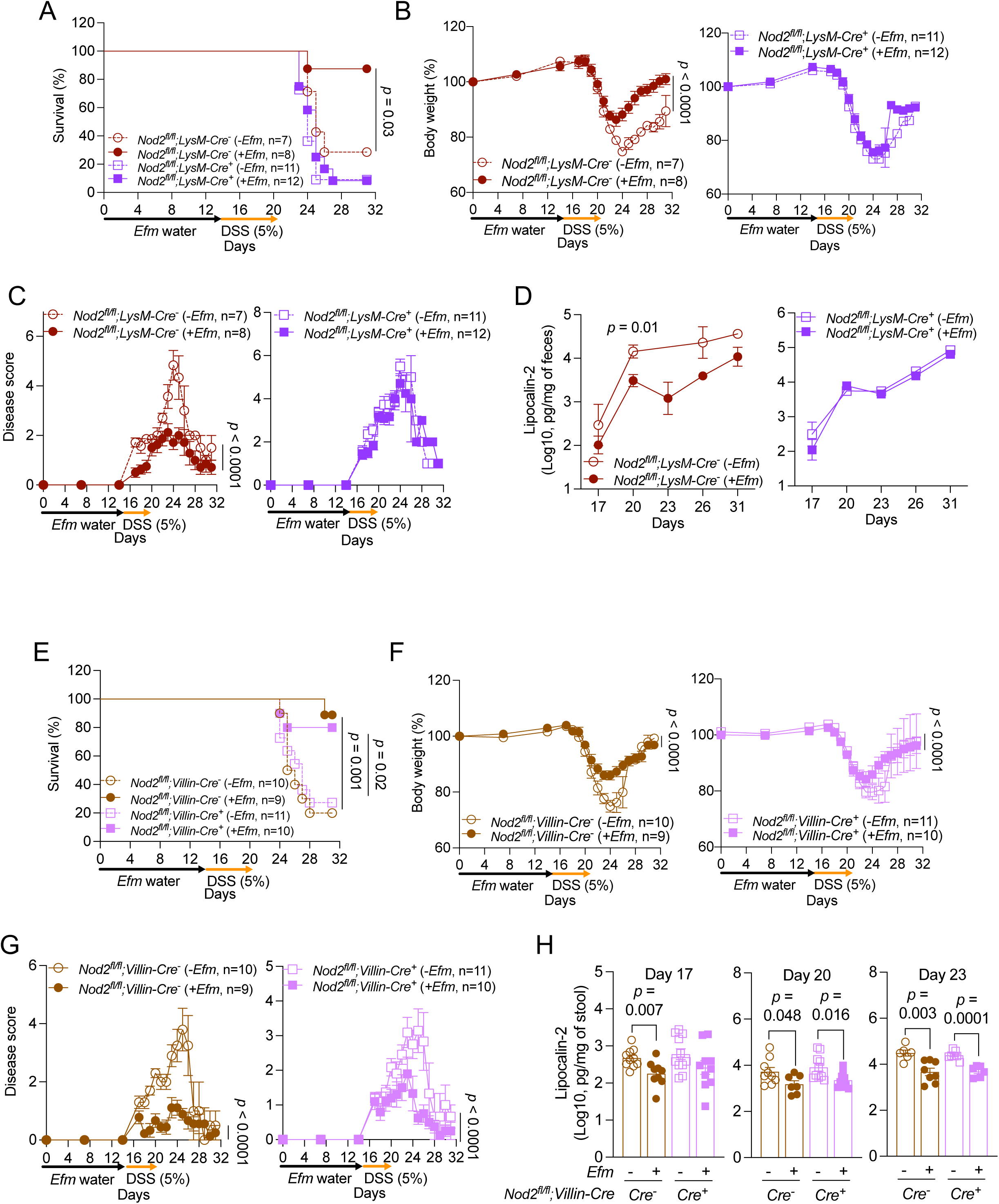
NOD2 in myeloid cells is required for *Efm*-mediated protection. **A-H)** DSS-treatment of *Nod2^fl/fl^;LysM-Cre*^-^ and *Cre*^+^ mice (A-D) or *Nod2^fl/fl^;Villin-Cre*^-^ and *Cre*^+^ mice (E-H) from room 6 following administration of *Efm* in drinking water or control. The mice were examined for survival (A and E), changes in body weight (B and F) and disease score (C and G), and LCN2 presence in stool samples at the indicated time points (D and H). Data points in H represent Individual mice. Data points in B-D, F, and G represent mean ± SEM. Bars represent mean ± SEM and at least three independent experiments were performed. Indicated *p* values by log-rank Mantel-Cox test in A and E, two-way ANOVA test in B, C, F, and G, and unpaired *t* test, two-tailed in D and H.

### *Efm* induces IL-22 production by lymphoid cells downstream of myeloid NOD2 to protect against intestinal injury

Activation of NOD2 in myeloid cells is frequently associated with pro-inflammatory cytokine production^63–65^, which is counterintuitive when considering the anti-inflammatory function observed in the above experiments. One way to reconcile these observations would be if myeloid cells, following NOD2 activation, act on other cell types in the gut to produce factors involved in tissue repair. For example, IL-22 is a major tissue regenerative cytokine and is protective in models of DSS-induced intestinal inflammation similar to the one used here^66, 67^. We observed increased IL-22 production in colon explants harvested from WT mice on day 14 following *Efm* administration in the drinking water (prior to DSS) and day 20 (with DSS) compared with mice that did not receive *Efm* (Figure 6A and B). IL-22 secretion was not enhanced by *Efm* colonization in *Nod2^-/-^* and *Nod2^fl/fl^;LysM-Cre^+^* mice on days 14 and 20 (Figure 6A and B), indicating that *Efm*-mediated induction of IL-22 is dependent on myeloid *Nod2* expression.

**Figure 6.**
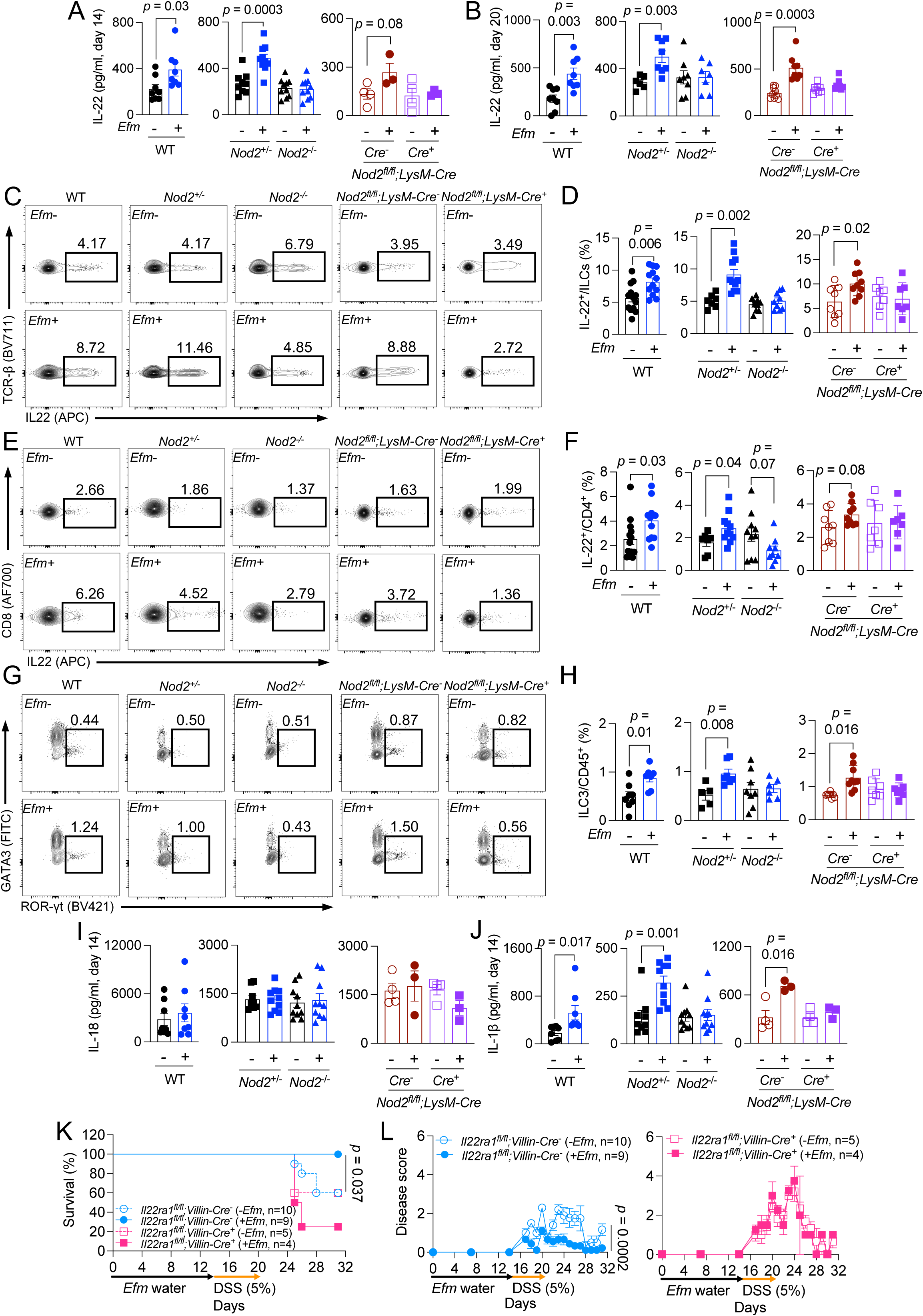
*Efm* induces IL-1β and increases the proportion of IL-22-producing lymphoid cells to prevent intestinal injury. **A and B)** Quantification of IL-22 on days 14 (A) and 20 (B) in the supernatants of gut explants from WT, *Nod2*^+/-^, *Nod2*^-/-^, and *Nod2^fl/fl^;LysM-Cre*^-^ and *Cre*^+^ mice with and without *Efm* inoculation. C-F) Representative flow cytometry plots and quantification of proportion of IL-22^+^ innate lymphoid cells (ILCs) (C and D) and CD4^+^ T helper cells (E and F) of WT, *Nod2*^+/-^, *Nod2*^-/-^, and *Nod2^fl/fl^;LysM-Cre*^-^ and *Cre*^+^ mice with and without *Efm* inoculation. G and H) Representative flow cytometry plots and quantification of group 3 ILCs (ILC3s) from mice in C-F. I and J) Quantification of IL-18 (I) and IL-1β (J) in the supernatants of gut explants from A. K and L) DSS treatment of *Il22ra1^fl/fl^;Villin-Cre^-^* and *Cre^+^* mice from room 6 following 2-week administration of *Efm* in drinking water or control. The mice were examined for survival (K) and changes in disease score (L). Data points in A, B, D, F, H, I, and J represent individual mice. For A, I, and J, n = 8 (WT), 8 (WT *Efm*), 9 (*Nod2*^+/-^), 10 (*Nod2*^+/-^ *Efm*), 10 (*Nod2^-/-^*), 10 (*Nod2*^-/-^ *Efm*), 4 (*Nod2^fl/fl^;LysM-Cre*^-^), 3 (*Nod2^fl/fl^;LysM-Cre*^-^ *Efm*), 4 (*Nod2^fl/fl^;LysM-Cre*^+^), and 3 (*Nod2^fl/fl^;LysM-Cre*^+^ *Efm*). For B, n = 8 (WT), 8 (WT *Efm*), 6 (*Nod2*^+/-^), 8 (*Nod2*^+/-^ *Efm*), 8 (*Nod2^-/-^*), 7 (*Nod2*^-/-^ *Efm*), 8 (*Nod2^fl/fl^;LysM-Cre*^-^), 8 (*Nod2^fl/fl^;LysM-Cre*^-^ *Efm*), 9 (*Nod2^fl/fl^;LysM-Cre*^+^), and 8 (*Nod2^fl/fl^;LysM-Cre*^+^ *Efm*). For D and F, n =13 (WT), 12 (WT *Efm*), 7 (*Nod2*^+/-^), 11 (*Nod2*^+/-^ *Efm*), 10 (*Nod2^-/-^*), 9 (*Nod2*^-/-^ *Efm*), 7 (*Nod2^fl/fl^;LysM-Cre*^-^), 7 (*Nod2^fl/fl^;LysM-Cre*^-^ *Efm*), 8 (*Nod2^fl/fl^;LysM-Cre*^+^), and 9 (*Nod2^fl/fl^;LysM-Cre*^+^ *Efm*). For H, n = 8 (WT), 7 (WT *Efm*), 5 (*Nod2*^+/-^), 7 (*Nod2*^+/-^ *Efm*), 8 (*Nod2^-/-^*), 6 (*Nod2*^-/-^ *Efm*), 7 (*Nod2^fl/fl^;LysM-Cre*^-^), 7 (*Nod2^fl/fl^;LysM-Cre*^-^ *Efm*), 8 (*Nod2^fl/fl^;LysM-Cre*^+^), and 9 (*Nod2^fl/fl^;LysM-Cre*^+^ *Efm*). Data points in L represent mean ± SEM. Bars represent mean ± SEM and at least two independent experiments were performed. Indicated *p* values by unpaired *t* test, two-tailed in A, B, D, F, H, and J, log-rank Mantel-Cox test in K, and two-way ANOVA test in L.

IL-22 in the gut is typically produced by group 3 innate lymphoid cells (ILC3s) and CD4^+^ T cells. To identify the subset of immune cells that induce IL-22 secretion responding to *Efm* colonization, we isolated colonic lamina propria of WT mice inoculated with *Efm* on day 14 to analyze the immune cell compartment by flow cytometry. Bacterial colonization increased the proportion of both IL-22^+^ ILCs and CD4^+^ T cells in WT mice, and not *Nod2^-/-^* and *Nod2^fl/fl^;LysM-Cre^+^* mice (Figure 6C-F). NOD2 in antigen-presenting cells can induce IL-10 production from regulatory T cells (Tregs)^68^. However, *Efm* did not alter the proportion of IL10^+^ CD4^+^ T cells, CD4^+^ T helper 1 (Tbet^+^) and 2 (GATA3^+^) cells, and Tregs (Supplemental figure 6C-F). Also, consistent with ILC3s being a source of IL-22, *Efm* colonization increased the proportion of ILC3s and not ILC1s and ILC2s in a NOD2-dependent manner (Figure 6G and H and Supplemental figure 6G and H). Therefore, *Efm* induces expansion of IL-22-producing CD4^+^ T cells and ILCs in a myeloid NOD2-dependent manner.

Secretion of IL-1β from intestinal myeloid cells induces IL-22 production from ILC3s^69, 70^. Also, myeloid cell-derived IL-1β secretion in the gut in response to the microbiota depends on NOD2^71^. The secretion of IL-1β, but not the related cytokine IL-18, was induced in *Efm*-colonized WT, *Nod2^+/-^*, and *Nod2^fl/fl^;LysM-Cre^-^* mice on days 14 and 20 (Figure 6I and J and Supplemental figure 6I and J). This *Efm*-mediated IL-1β induction did not occur in *Nod2^-/-^* and *Nod2^fl/fl^;LysM-Cre^+^* mice, indicating that *Nod2*-dependent sensing of *Efm* in myeloid cells is required for the upregulated IL-1β secretion (Figure 6I and J and Supplemental figure 6I and J). Therefore, NOD2 in myeloid cells mediates IL-1β secretion in response to *Efm* colonization.

To confirm whether IL-22 is required for *Efm*-mediated protection, we developed *Il22ra1^fl/fl^;Villin-Cre* mice in which *Il22ra1* encoding IL-22 receptor subunit α is deficient in intestinal epithelial cells and confirmed the *Efm* shedding in stool samples following inoculation in all groups (Supplemental figure 6K). *Il22ra1^fl/fl^;Villin-Cre^+^* mice displayed exacerbated signs of intestinal injury compared with *Il22ra1^fl/fl^;Villin-Cre^-^* mice regardless of *Efm* colonization (Figure 6K and L and Supplemental figure 6L). In contrast, *Efm* colonization improves mortality, body weight reduction, and disease score in *Il22ra1^fl/fl^;Villin-Cre^-^* mice (Figure 6K and L and Supplemental figure 6L). Therefore, epithelial response to IL-22 is necessary for *Efm*-mediated protection.

### The NOD2 R702W equivalent impairs *Efm*-mediated protection in mice

The three major NOD2 variants linked to IBD − R702W, G908R, and a frameshift deletion mutation at L1007 (L1007fs) − result in the loss of muropeptide recognition and NF-κB signaling *in vitro*^72, 73^. L1007fs has received the bulk of attention, and mice harboring a similar frameshift mutation display a compromised cytokine response during bloodstream infection by *Efl*^42^, confirming that it results in loss-of-function. The R702W variant, despite being the most common and detected in up to 5% of individuals of European descent (Supplemental figure 7A), has not been studied using an *in vivo* model (Figure 7A). Thus, we introduced a knock-in mutation to generate mice harboring the equivalent of the human R702W variant (Q675W, Supplemental figure 7B). We confirmed the absence of other mutations in *Nod2,* and that NOD2 Q675W protein was produced (Supplemental figure 7C and D). After backcrossing with WT B6 mice, we used heterozygous mice (*Nod2^Q^*^675^*^W/+^*) as littermate controls. Like other mice, *Efm* was detected in stool collected from *Nod2^Q^*^675^*^W/+^* and *Nod2^Q^*^675^*^W/Q675W^* following administration in drinking water (Supplemental figure 7E). In the absence of *Efm*, *Nod2^Q675W/+^* and *Nod2^Q675W/Q675W^* mice displayed modest mortality, weight loss, or disease score following DSS treatment (Figure 7B-D), likely due to the high levels of endogenous *Enterococcus* colonization at the outset of the experiment compared with the other mice raised in the same room (Supplemental figure 7G). Nevertheless, we were able to detect significant differences between *Nod2^Q675W/+^* and *Nod2^Q675W/Q675W^* mice. *Efm* colonization did not improve survival, body weight reduction, disease score, fecal LCN2 concentration, colon shortening, and intestinal histopathology in *Nod2^Q675W/Q675W^* mice, and as such, *Efm* colonized mice were more susceptible to DSS than similarly treated *Nod2^Q675W/+^* mice (Figure 7B-H, Supplemental figure 7F, and Supplemental table 3). These results indicate that the NOD2 R702W equivalent in mice impairs *Efm*-mediated protection.

**Figure 7.**
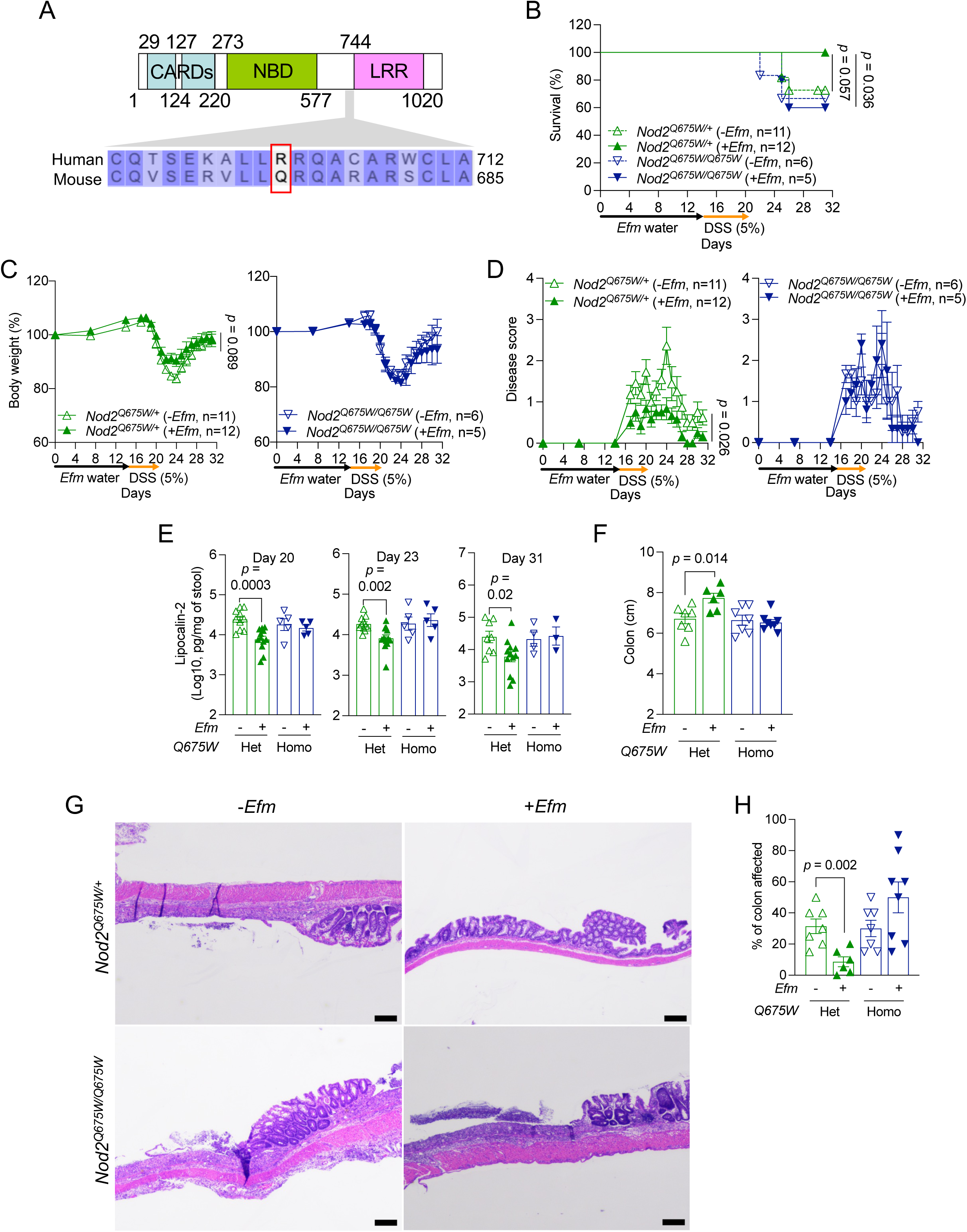
NOD2 Q675W impairs *Efm*-mediated protection. **A)** NOD2 protein region targeted by mutagenesis. Arginine at position 702 in human NOD2 and the corresponding glutamine at position 675 in mouse NOD2 are indicated in the red box. Numbers refer to human amino acid positions. **B-F)** DSS-treated *Nod2^Q675W/+^* and *Nod2^Q675W/Q675W^* mice with and without *Efm* inoculation were examined for survival (B), changes in body weight (C) and disease score (D), and LCN2 presence in stool samples at the indicated time points (E), and colon length (F) on day 23. **G and H)** Representative images of H&E-stained sections of the colon (G) and quantification of the proportion of colon affected (H). Bars, 200 μM. Data points in E, F, and H represent individual mice. Data points in C and D represent mean ± SEM. Bars represent mean ± SEM and at least two independent experiments were performed. CARD, caspase recruitment domain; NBD, nucleotide-binding domain; LRR, leucine-rich repeat domain; Het, heterozygotes; Homo, homozygotes. Indicated *p* values by log-rank Mantel-Cox test in B, two-way ANOVA test in C and D, and unpaired *t* test, two-tailed in E, F, and H.

## DISCUSSION

In patients with IBD, previous small studies demonstrated excessive *REG3* gene expression despite that AMP secretion is generally associated with an improved mucosal barrier^39^. Here, we revisited this unresolved counterintuitive observation. Functional assays showed increased REG3 antimicrobial activity in IBD patients compared with controls matched for gastrointestinal symptoms, indicating that REG3 overproduction is not a general byproduct of diarrheal disease and more likely IBD itself. Based on prior studies demonstrating that enterococci are among the most sensitive bacteria to REG3-mediated killing^6, 9^, we focused our analysis on this group and showed their reduced presence in IBD patients. Recent findings indicating that *Enterococcus* species activate the IBD susceptibility gene NOD2 led us to hypothesize that these bacteria have anti-inflammatory properties, which we tested in an animal model of intestinal injury. The results from these experiments support a model in which SagA-secreting enterococci such as *Efm* activate NOD2 in myeloid cells to induce IL-1β, leading to an increase in lymphoid cell types that produce the protective cytokine IL-22. Once the inflammation has been resolved, this enhanced IL-22 may maintain optimal REG3 expressions through STAT3 activation^74^, perpetuating gut homeostasis. In IBD, either depletion of enterococci due to REG3 overproduction or genetic deficiency in NOD2 function disrupts this anti-inflammatory circuit (Supplemental figure 7H). This model helps explain how IBD patients with intact NOD2 can develop disease presentations similar to those observed in individuals with NOD2 variants once inflammation is established.

Excess REG3 likely acts in concert with other mechanisms to disrupt the microbiota of IBD patients. Altered availability of oxygen and nitrate in the inflamed gut favors colonization by facultative anaerobic Enterobacteriaceae species over the strictly anaerobic *Firmicutes*, the phylum that includes enterococci^75, 76^. A recent study identified additional *Firmicutes* that encode NOD2-activating DL-endopeptidases such as *Streptococcus cristatus*, *Lactobacillus rhamnosus*, and *Bacteroides caccae*, which are reduced in IBD patients^40^. It is possible that AMP dysregulation exacerbates the depletion of NOD2-activating species that are already vulnerable due to imbalanced redox status of the inflamed environment. Notably, antimicrobial activity was greater in the IBD cohort compared with individuals experiencing gastrointestinal symptoms without IBD. REG3 overproduction may explain why individuals who experience transient gastroenteritis recover while individuals with IBD remain susceptible to disease flare.

Our findings have relevance to other disease settings such as cancer. Probiotics engineered to secrete SagA were shown to improve the efficacy of immune checkpoint blockade therapy in reducing tumors in mice^50^. Although adverse events such as colitis have been a major concern for strategies that improve antitumor therapies, our results predict that inducing NOD2 through SagA would promote checkpoint inhibitor efficacy while also reducing intestinal inflammation. However, our findings with the *Nod2 Q675W* knock-in mice suggest that this approach will fail in individuals homozygous for analogous loss-of-function variants of NOD2. Genetic information may be useful in determining eligibility of patients for microbiota-targeting therapies. Additionally, in treating various diseases, administration of live bacteria is associated with substantial safety concerns. In this context, it is notable that we achieved a similar degree of protection against intestinal injury by administering sterile supernatant collected from SagA-producing bacterial cultures. Postbiotic strategies using bacterial products may be a safe alternate to beneficially trigger NOD2.

### Limitations of the study

The consequence of *Enterococcus* depletion on intestinal inflammation and the immune mechanism was investigated in an animal model. A long-term goal would be to determine whether NOD2 activation by enterococci leads to similar immune responses in humans and ultimately tests the efficacy of therapies designed based on such findings. However, there are significant safety concerns when considering interventions that target these pathways. For instance, IL-1β production promotes intestinal inflammation in the IL-10-deficient setting^77^. Also, our proposed immune circuit induced by NOD2 in myeloid cells is not mutually exclusive with the multitude of other ways in which NOD2 has been shown to protect the intestinal barrier.

## Figure legend

**Supplemental figure 1.**
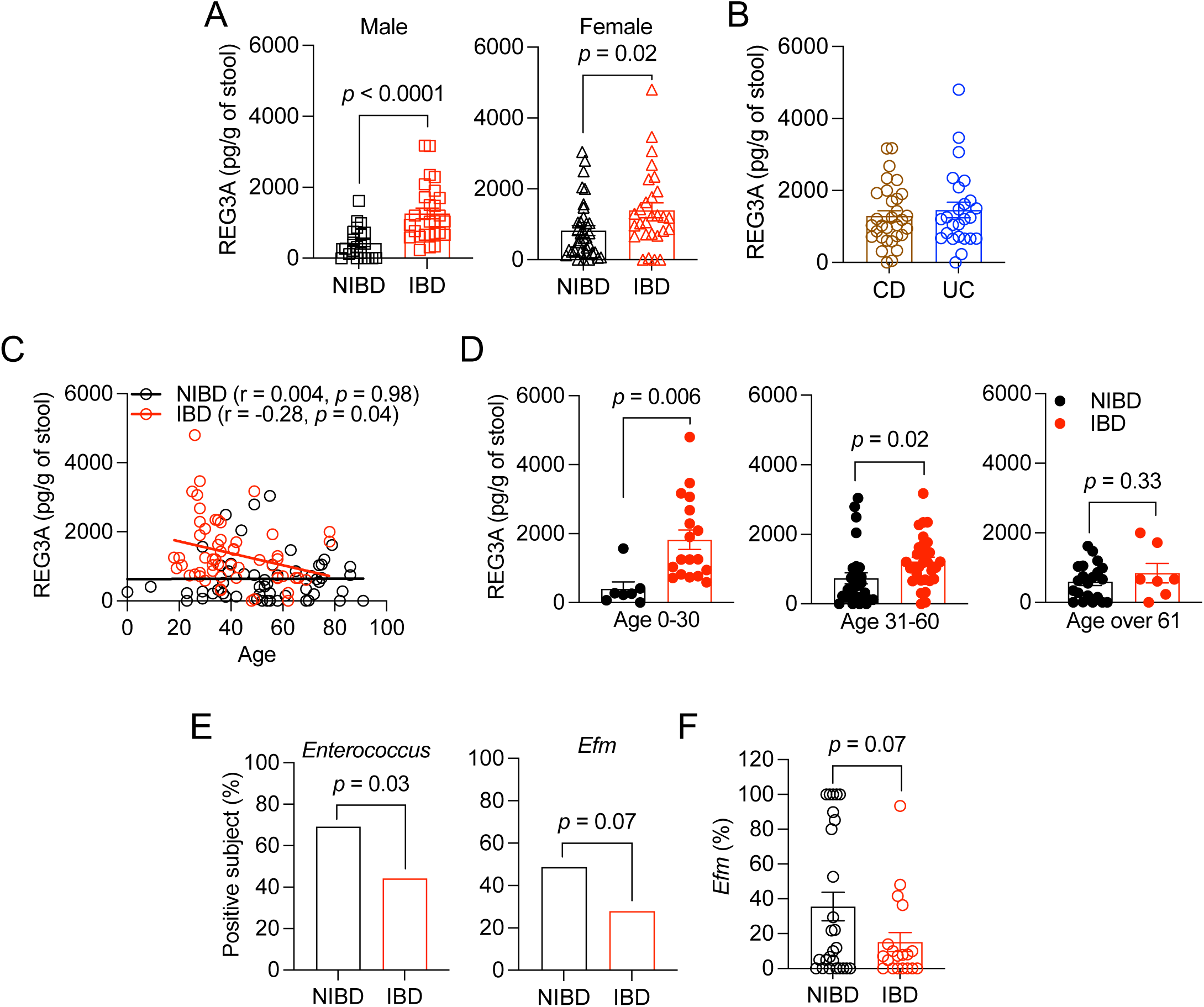
Contribution of patient demographics to REG3A levels and antimicrobial activity. **A)** REG3A concentration in stool extracts from NIBD and IBD patients from Figure 1 separated by sex. **B)** REG3A concentration in stool extracts from Crohn’s disease (CD) or ulcerative colitis (UC) patients. **C)** Correlation between age of NIBD and IBD patients and REG3A concentration. **D)** REG3A concentration in stool extracts segregated into indicated age groups. **E)** Proportion of patients in which *Enterococcus* and *E. faecium* (*Efm*) were detected in stool. **F)** Proportion of *Enterococcus* that are *Efm* in NIBD and IBD stools. Data points in A-D and F represent individual patients. Bars represent mean ± SEM and at least two independent experiments were performed. r, Pearson correlation coefficient. Indicated *p* values by unpaired *t* test, two-tailed in A, D, and F, simple linear regression analysis in C, and Fisher’s exact test in E.

**Supplemental figure 2.**
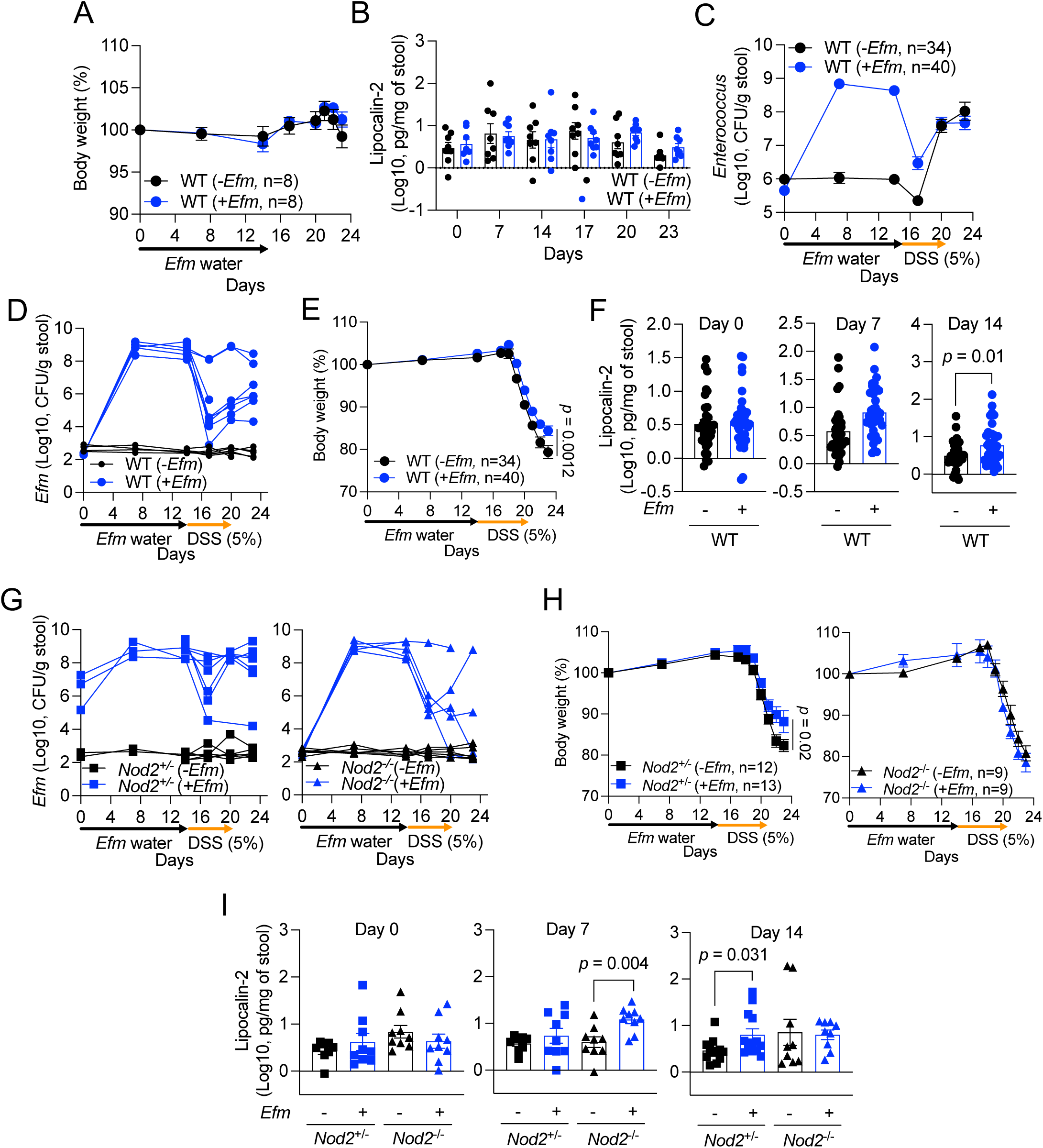
*Efm* colonization levels and disease parameters in wild-type and *Nod2*-deficient mice. **A and B)** Changes in body weight (A) and lipocalin-2 (LCN2) concentration in stool samples at the indicated time points (B) in wild-type (WT) B6 mice following administration of *Efm* in drinking water or control. **C-F)** Dextran sulfate sodium (DSS)-induced intestinal injury of WT mice from room 6 following administration of *Efm* in drinking water or control. The mice were examined for the burden of total *Enterococcus* (C) and *Efm* (D) in the stool, changes in body weight (E), and fecal LCN2 concentration (F) at the indicated time points. **G-I)** DSS-treatment of *Nod2*^+/-^ and *Nod2*^-/-^ mice from room 6 following administration of *Efm* in drinking water or control. The mice were examined for the burden of *Efm* (G) in the stools, changes in body weight (H), and fecal LCN2 concentration (I) at the indicated time points. Data points in B, F, and I and lines in D and G represent individual mice. Data points in A, C, E, and H represent mean ± SEM. Bars represent mean ± SEM and at least three independent experiments were performed. Indicated *p* values by two-way ANOVA test in E and H and unpaired *t* test, two-tailed in F and I.

**Supplemental figure 3.**
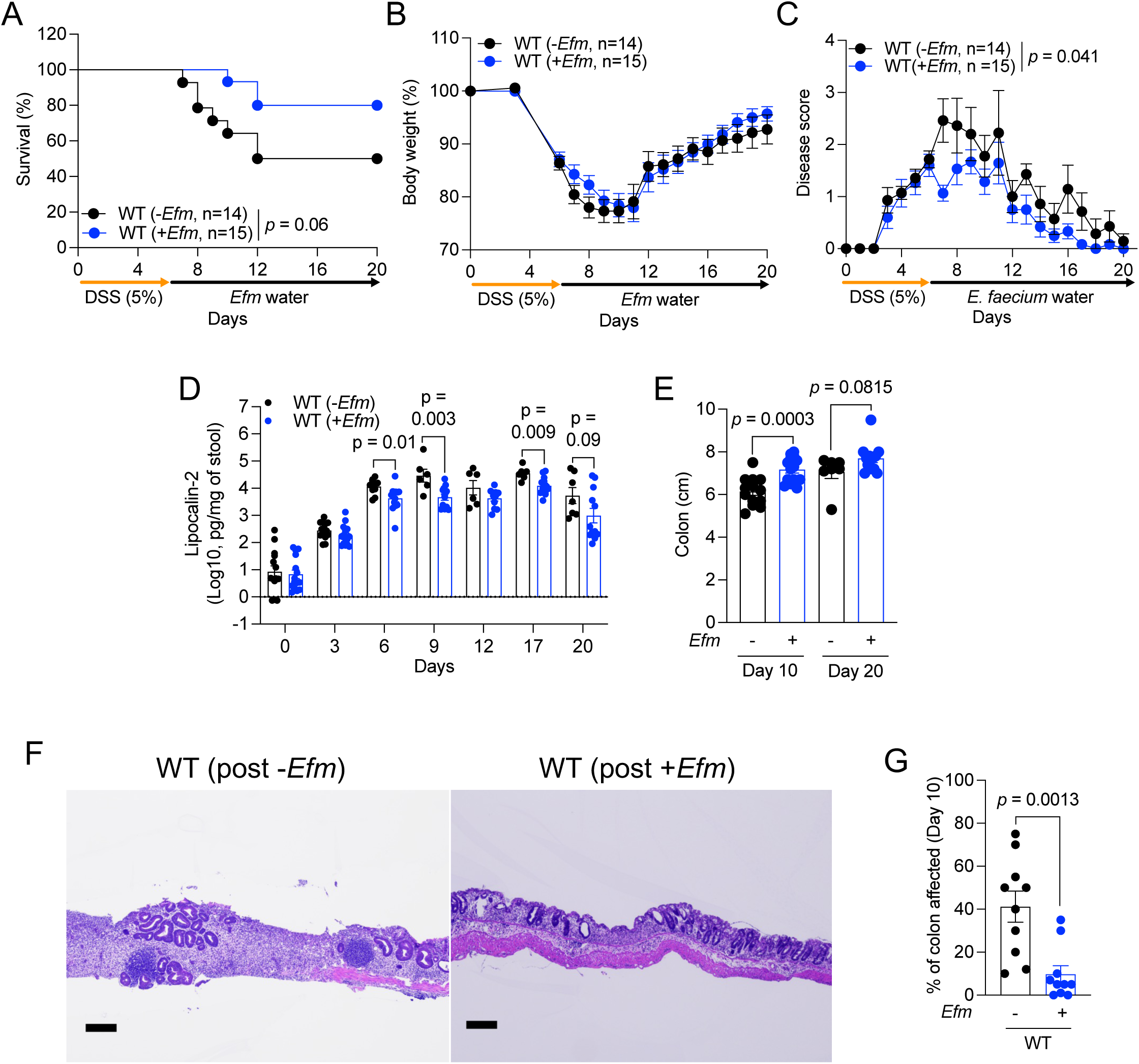
*Efm* inoculation after DSS administration ameliorates intestinal injury. **A-E)** WT B6 mice from room 6 received *Efm* in drinking water or control following DSS treatment. The mice were examined for survival (A), changes in body weight (B) and disease score (C), and LCN2 concentration (D) in stool samples at the indicated time points, colon length on days 10 and 20 (E). **F and G)** Representative images of hematoxylin and eosin (H&E)-stained sections of the colon (F) and quantification of the proportion of colon affected (G) on day 10 from mice in A-E. Bars, 200 μM. Data points in D, E, and G represent individual mice from mice in A-C. Data points in B and C represent mean ± SEM. Bars represent mean ± SEM and at least three independent experiments were performed. Indicated *p* values by log-rank Mantel-Cox test in A, two-way ANOVA test in C, unpaired *t* test, two-tailed in D, E, and G.

**Supplemental figure 4.**
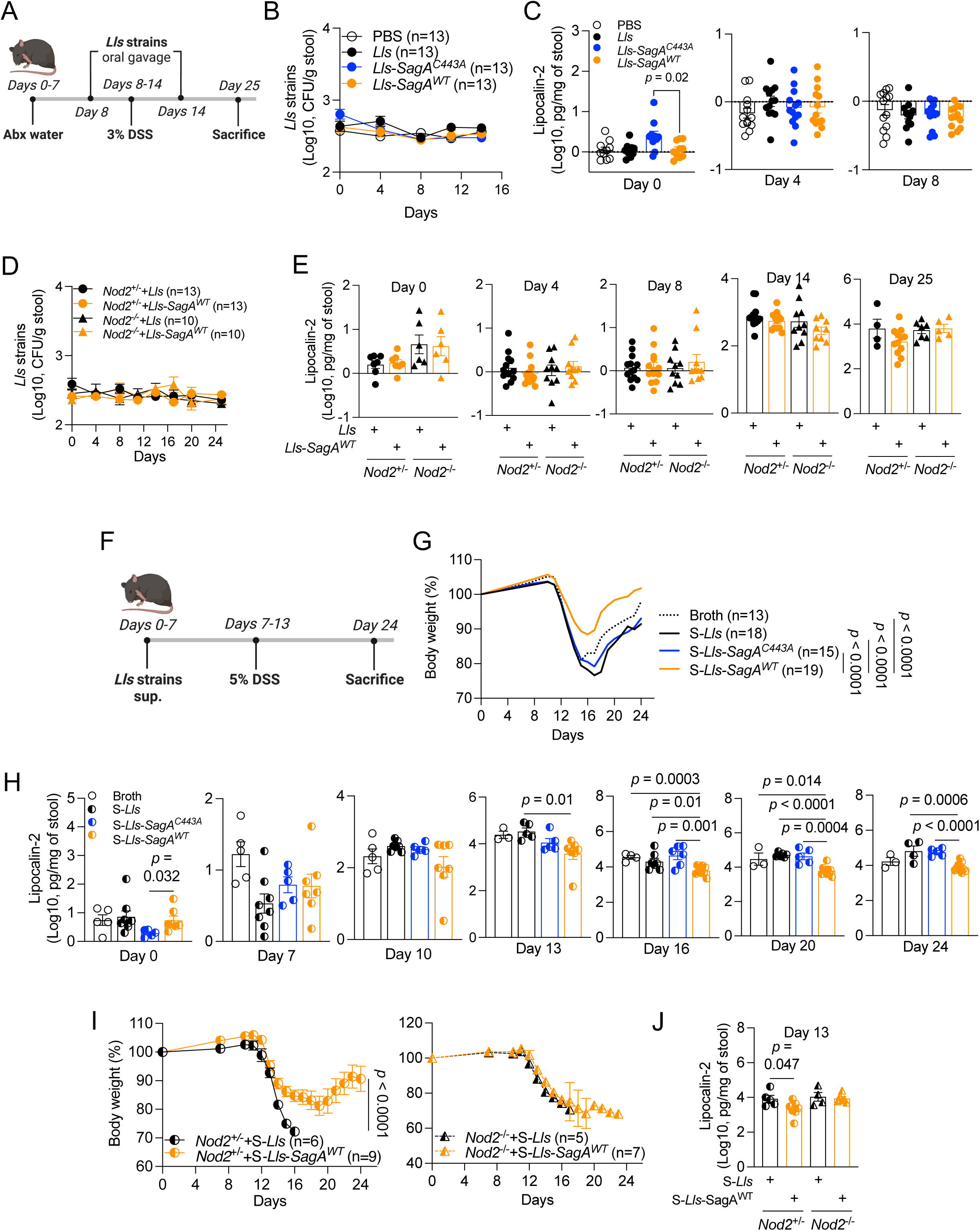
*L. lactis* expressing SagA promotes NOD2-dependent protection against intestinal injury. **A)** Schematic of experimental procedure for mice inoculation with *L. lactis* (*Lls*) strains. Mice received antibiotics (ampicillin + streptomycin, abx) in drinking water for 7 days, and were orally gavaged with PBS or *Lls* strains twice on days 8 and 14 to coincide with the initiation and cessation of DSS administration. **B and C)** Abx-treated WT mice given PBS, parental *Lls*, *Lls* expressing catalytic mutant SagA (*Lls*-*SagA^C443A^*), or *Lls* expressing WT SagA (*Lls*-*SagA^WT^*) were treated with DSS as in A and examined for the burden of *Lls* strains (B) and LCN2 concentration in stool samples at the indicated time points (C). **D and E)** Abx-treated *Nod2*^+/-^ and *Nod2*^-/-^ mice given parental *Lls* or *Lls*-*SagA^WT^* were treated with DSS and examined for the burden of *Lls* strains (D) and LCN2 concentration in stool samples at the indicated time points (E). **F)** Schematic of experimental procedure for DSS treatment of mice that received broth or culture supernatants of *Lls* strains as drinking water. **G and H)** DSS-treatment of WT mice inoculated with broth or culture supernatants of *Lls* (S-*Lls*), *Lls*-*SagA^C443A^* (S-*Lls*-*SagA^C443A^*), or *Lls*-*SagA^WT^* (S-*Lls*-*SagA^WT^*) were examined for changes in body weight (G) and LCN2 presence in stool samples at the indicated time points (H). **I and J)** DSS-treatment of *Nod2*^+/-^ and *Nod2*^-/-^ mice inoculated with S-*Lls* or S-*Lls*-*SagA^WT^*. The mice were examined for changes in body weight (I) and LCN2 presence in stool samples on day 13 (J). All mice were bred in room 6. Data points in C, E, H, and J represent individual mice. Data points in B, D, and I represent mean ± SEM. Lines in G represent mean. Bars represent mean ± SEM and at least two independent experiments were performed. Indicated *p* values by unpaired *t* test, two-tailed in C, H, and J and two-way ANOVA test in G and I.

**Supplemental figure 5.**
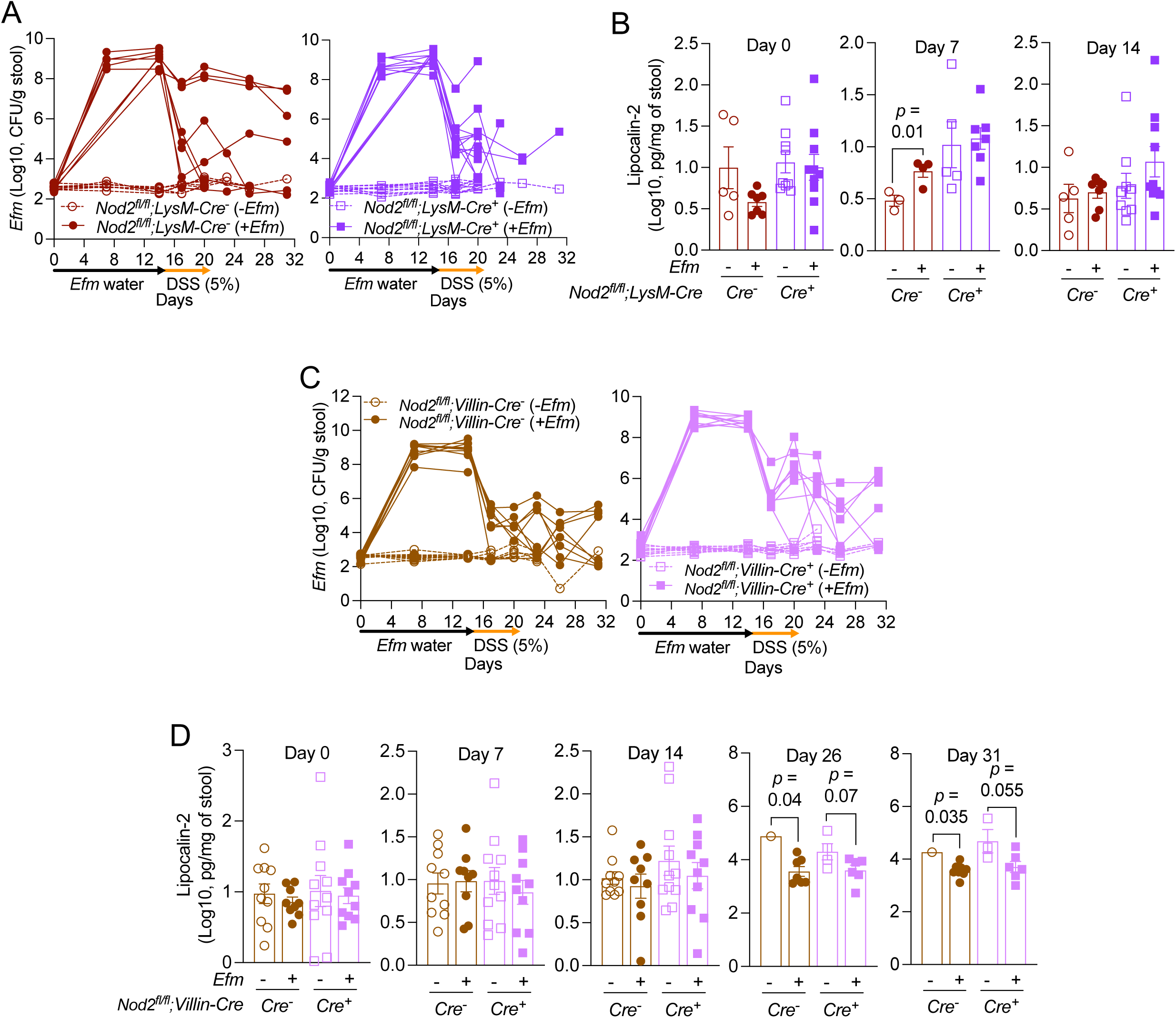
*Efm*-mediated protection is dependent on myeloid and not intestinal epithelial *Nod2* expression. **A-D)** DSS-treatment of *Nod2^fl/fl^;LysM-Cre*^-^ and *Cre*^+^ mice (A and B) or *Nod2^fl/fl^;Villin-Cre*^-^ and *Cre*^+^ mice (C and D) from room 6 following administration of *Efm* in drinking water or control. The mice were examined for *Efm* burden in stool samples (A and and fecal LCN2 concentration at the indicated time points (B and D). Lines in A and C and data points in B and D represent individual mice. Bars represent mean ± SEM and at least three independent experiments were performed. Indicated *p* values by unpaired *t* test, two-tailed in B and D.

**Supplemental figure 6.**
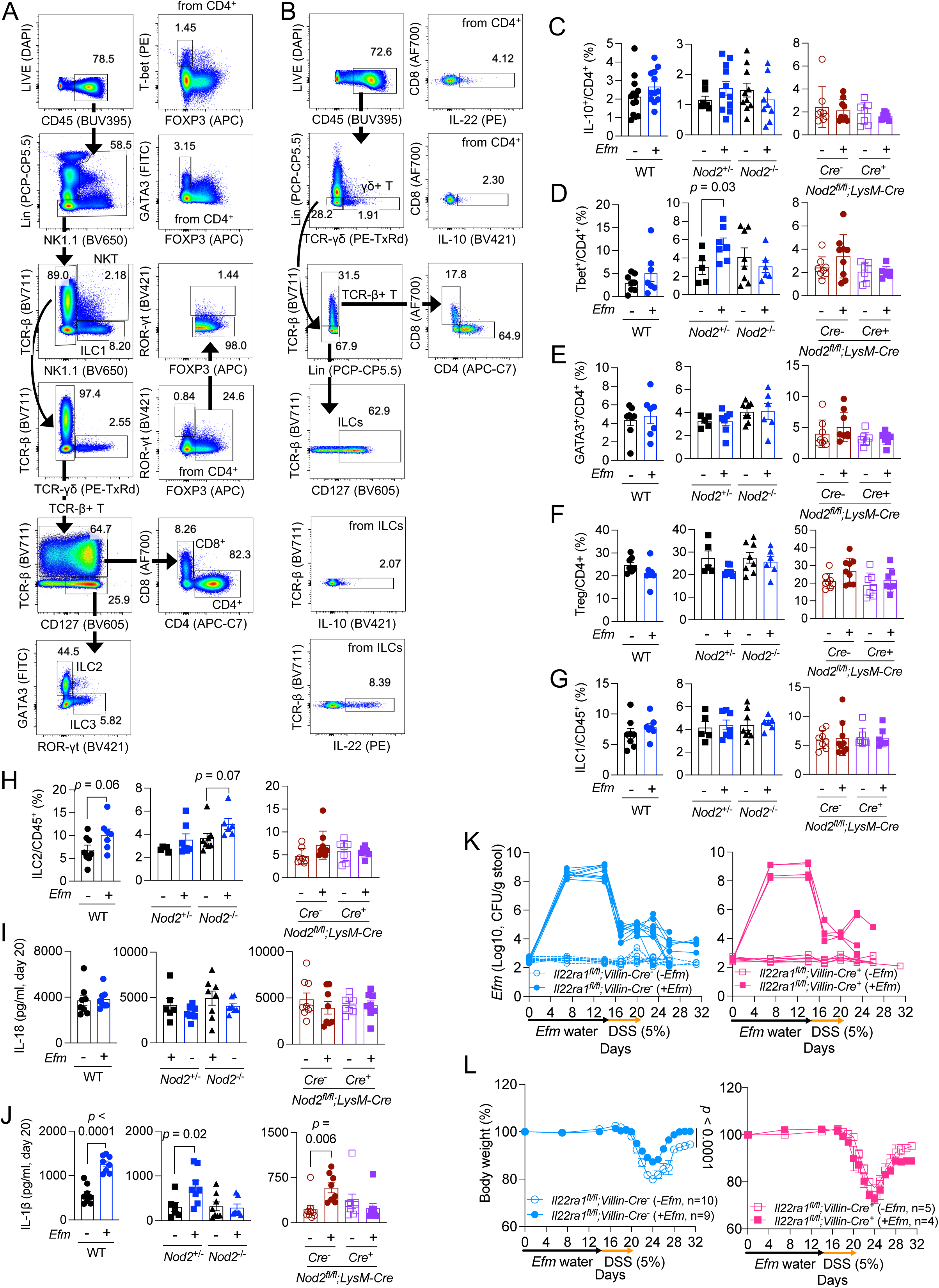
Immune cell and cytokine analysis of colons from *Efm*-colonized mice and analysis of *Efm*-colonized *Il22ra1^fl/fl^;Villin-Cre* mice. **A and B)** Flow cytometry gating strategies for T cells and innate lymphoid cells (ILCs) (A, transcription factors), and cytokine production (B). C-F) Flow cytometry graph depicting the proportion of IL10^+^ (C), Tbet^+^ (D), and GATA3^+^ (E) CD4^+^ T cells and regulatory T cells (Treg, F) in WT, *Nod2*^+/-^, *Nod2*^-/-^, and *Nod2^fl/fl^;LysM-Cre*^-^ and *Cre*^+^ mice following 2-week administration of *Efm* in drinking water or control. G and H) Flow cytometry graph depicting the proportion of group 1 (ILC1, G) and group 2 (ILC2, H) ILCs in WT, *Nod2*^+/-^, *Nod2*^-/-^, and *Nod2^fl/fl^;LysM-Cre*^-^ and *Cre*^+^ mice following 2-week administration of *Efm* in drinking water or control. I and J) Quantification of secreted IL-18 (I) and IL-1β (J) in the supernatants of gut explants from *Efm*-colonized WT, *Nod2*^+/-^, *Nod2*^-/-^, and *Nod2^fl/fl^;LysM-Cre*^-^ and *Cre*^+^ mice on day 20. K and L) DSS treatment of *Il22ra1^fl/fl^;Villin-Cre^-^* and *Cre^+^* mice from room 6 following 2-week administration of *Efm* in drinking water or control. The mice were examined for the burden of *Efm* (K) and changes in body weight (L). Data points in C-J and lines in K represent individual mice. For C, n = 13 (WT), 12 (WT *Efm*), 7 (*Nod2*^+/-^), 11 (*Nod2*^+/-^ *Efm*), 10 (*Nod2^-/-^*), 9 (*Nod2*^-/-^ *Efm*), 7 (*Nod2^fl/fl^;LysM-Cre*^-^), 7 (*Nod2^fl/fl^;LysM-Cre*^-^ *Efm*), 8 (*Nod2^fl/fl^;LysM-Cre*^+^), and 9 (*Nod2^fl/fl^;LysM-Cre*^+^ *Efm*). For D-H, n = 8 (WT), 7 (WT *Efm*), 5 (*Nod2*^+/-^), 7 (*Nod2*^+/-^ *Efm*), 8 (*Nod2^-/-^*), 6 (*Nod2*^-/-^ *Efm*), 7 (*Nod2^fl/fl^;LysM-Cre*^-^), 7 (*Nod2^fl/fl^;LysM-Cre*^-^ *Efm*), 8 (*Nod2^fl/fl^;LysM-Cre*^+^), and 9 (*Nod2^fl/fl^;LysM-Cre*^+^ *Efm*). For I and J, n = 8 (WT), 8 (WT *Efm*), 6 (*Nod2*^+/-^), 8 (*Nod2*^+/-^ *Efm*), 8 (*Nod2^-/-^*), 7 (*Nod2*^-/-^ *Efm*), 8 (*Nod2^fl/fl^;LysM-Cre*^-^), 8 (*Nod2^fl/fl^;LysM-Cre*^-^ *Efm*), 9 (*Nod2^fl/fl^;LysM-Cre*^+^), and 8 (*Nod2^fl/fl^;LysM-Cre*^+^ *Efm*). Data points in L represent mean ± SEM. Bars represent mean ± SEM and at least two independent experiments were performed. Indicated *p* values by unpaired *t* test, two-tailed in C-J and two-way ANOVA test in L.

**Supplemental figure 7.**
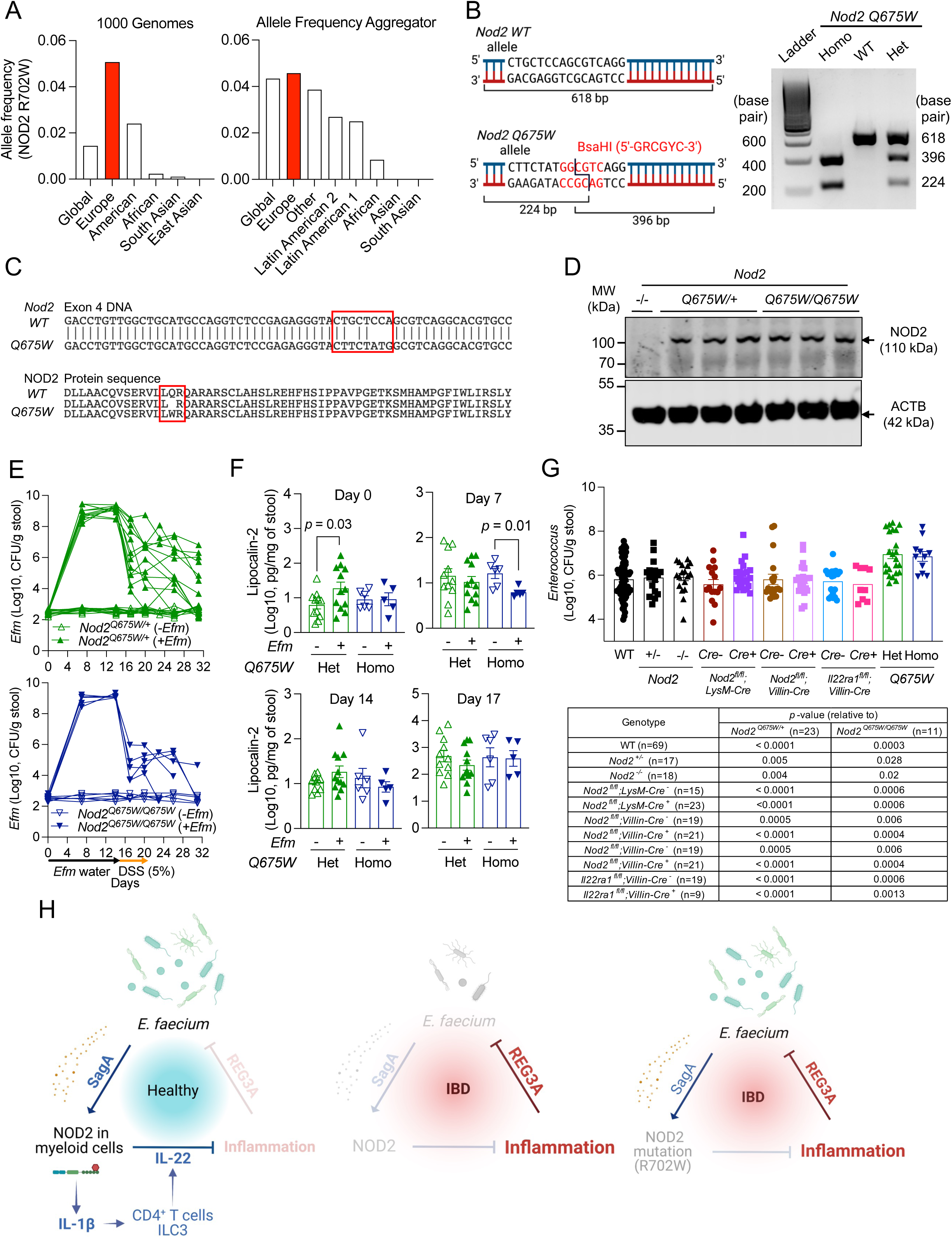
Characterization of mice carrying a *Nod2* knock-in mutation equivalent to the human R702W. **A)** *NOD2 R702W* allele frequency according to ethnic groups. The frequencies were retrieved from 1000 Genomes Project and Allele Frequency Aggregator. **B)** Schematic of genotyping PCR product (left) and representative genotyping gel image (right) for *Nod2 Q675W* knock-in mouse. **C)** Sequencing of the *Nod2* locus from the above mice confirmed successful gene targeting. Figure shows exon 4 DNA and amino acid sequences from *Nod2 Q675W* knock-in mouse sequencing results aligned to the WT sequences. Red boxes indicate the mutated region. **D)** Western blot image of NOD2 and β-actin (ACTB) in colonic tissue lysates from *Nod2*^-/-^, *Nod2^Q675W/+^*, and *Nod2^Q675W/Q675W^* mice. **E and F)** DSS treatment of *Nod2^Q675W/+^ and Nod2^Q675W/Q675W^* mice from room 6 following 2-week administration of *Efm* in drinking water or control. The mice were examined for the burden of *Efm* (E) and LCN2 concentration (F) in the stool samples at the indicated time points. **G)** Endogenous *Enterococcus* burden among mice with different genotypes on day 0 (upper). The *p*-values relative to *Nod2^Q675W/+^* or *Nod2^Q675W/Q675W^* mice were indicated in lower table. **H)** Schematic of the mechanism by which *Efm* activates NOD2 to suppress inflammation, and how this process is disrupted when either REG3A is overproduced or NOD2 is genetically inactivated. Lines in E and data points in F and G represent individual mice. Bars in F and G represent mean ± SEM and at least three independent experiments were performed. Het, heterozygotes; Homo, homozygotes. Indicated *p* values by unpaired *t* test, two-tailed in F and G.

## STAR METHODS

Detailed methods are provided in the online version of this paper and include the following:

- KEY RESOURCES TABLE
- RESOURCE AVAILABILITY

- Lead contact
- Materials availability
- Data and code availability
- EXPERIMENTAL MODEL AND SUBJECT DETAILS

- Mice
- Generation of *Nod2 Q675W* knock-in mice
- Human stool samples and data collection
- METHOD DETAILS

- Bacterial species
- Preparation of human stool extract, antimicrobial activity and REG3A measurement
- Western blotting
- DNA extraction and 16S rRNA sequencing analysis
- Enumeration of total *Enterococcus* and *Efm* in human stool samples
- Bacterial inoculation and DSS treatment of mice
- Quantification of bacterial burden and lipcalin-2 (LCN2) in murine stool samples
- Histology
- Colon explant culture and cytokine detection
- Isolation of lamina propria (LP) cells
- Flow cytometry
- *Nod2* sequencing
- Statistical analyses

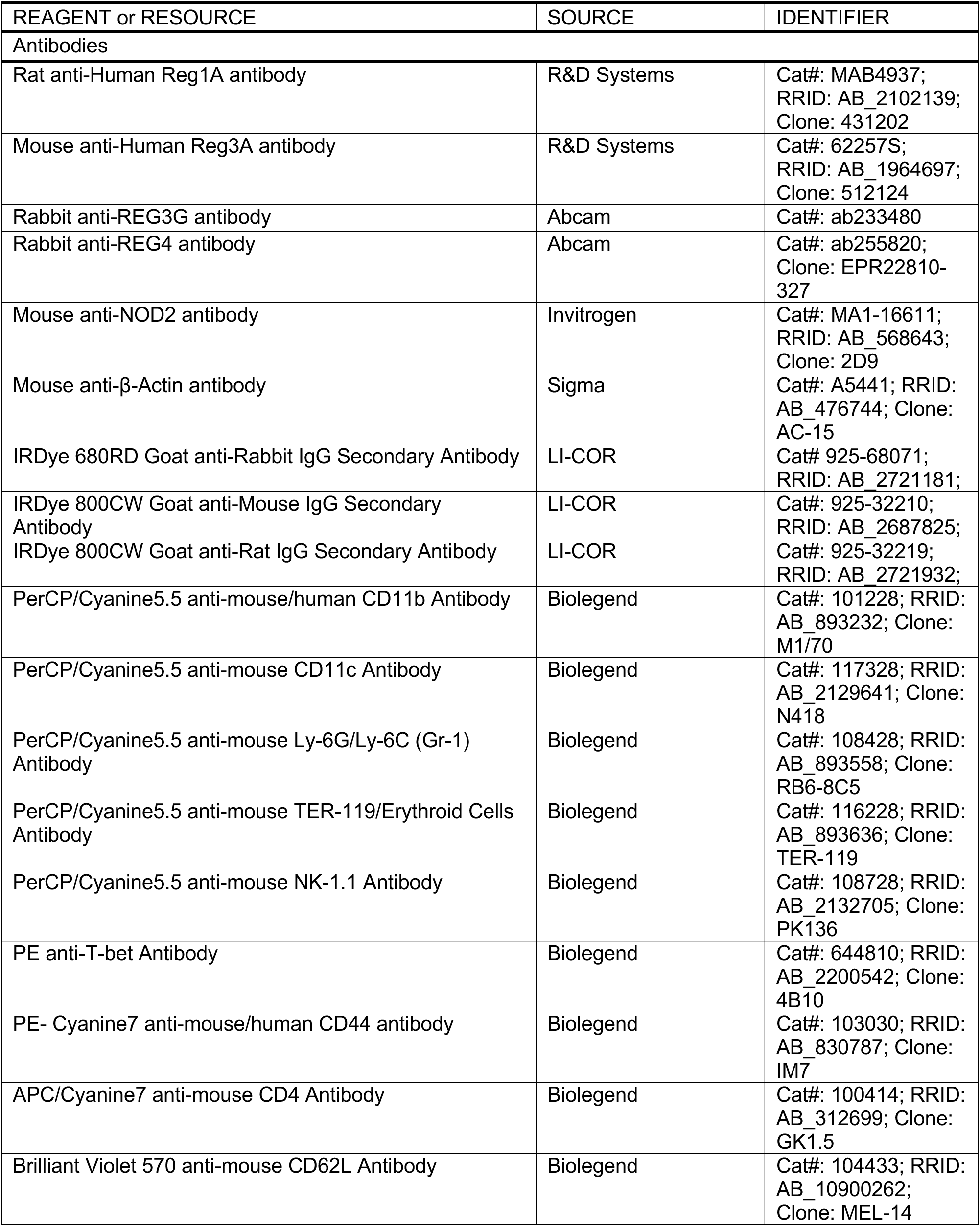
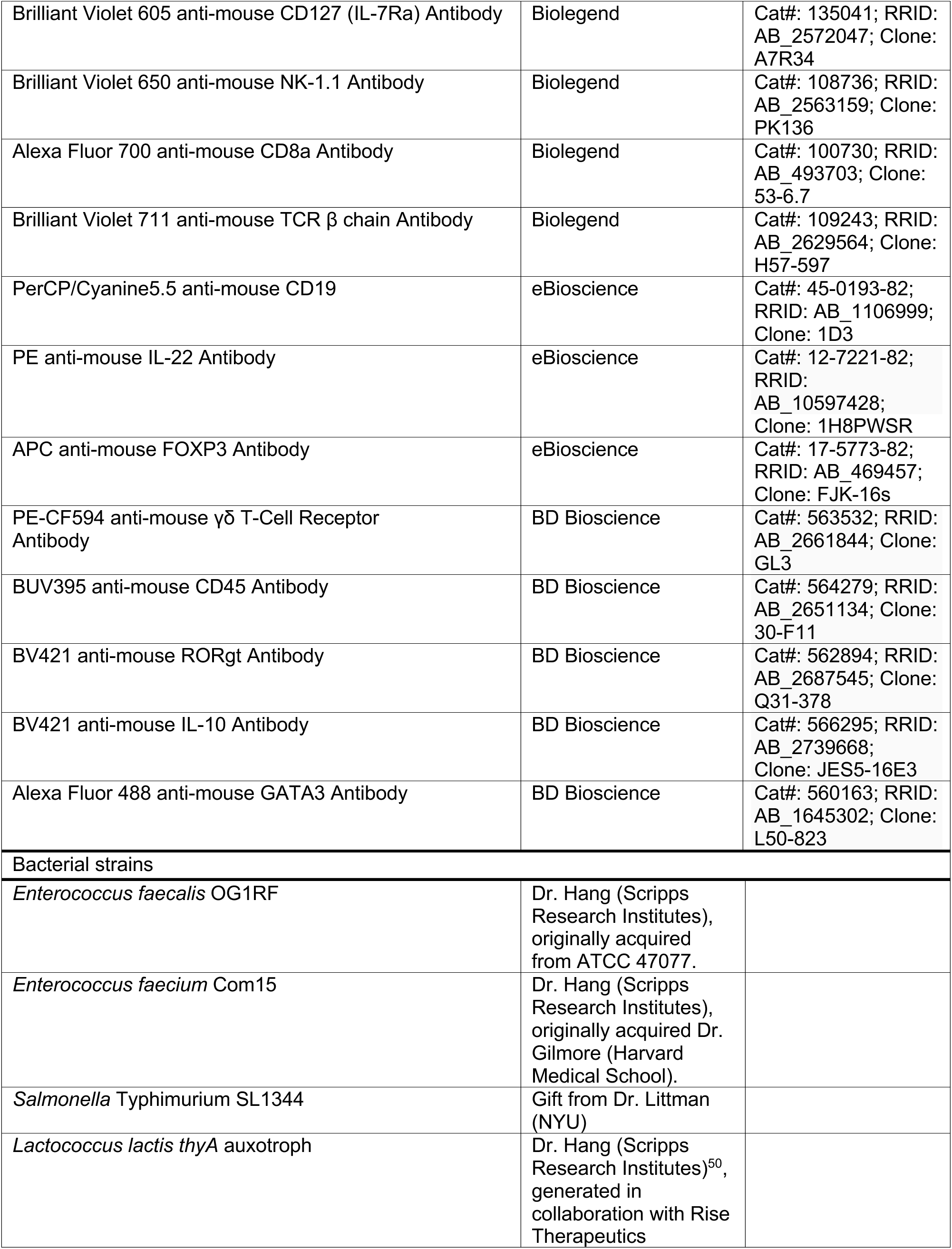
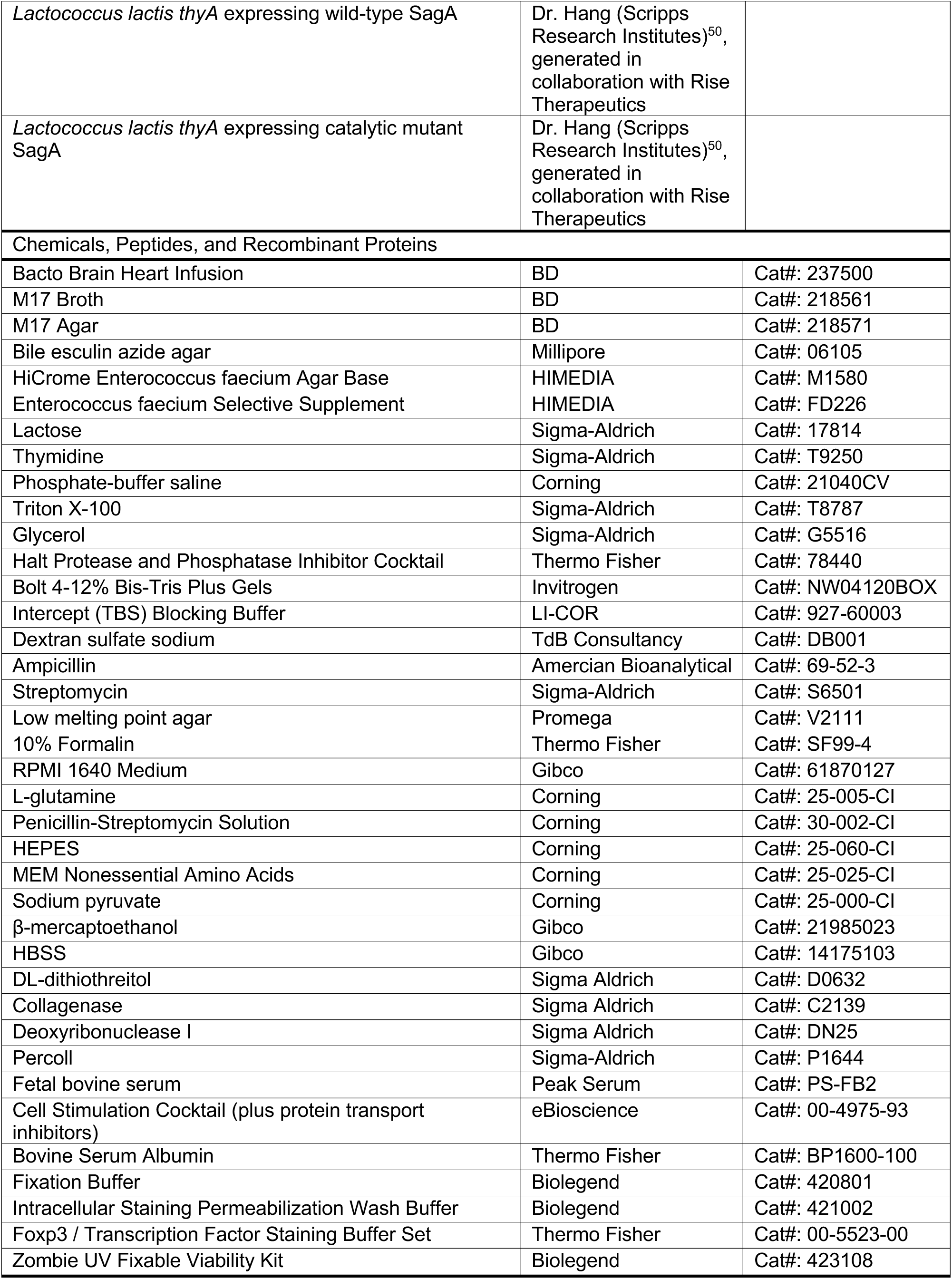
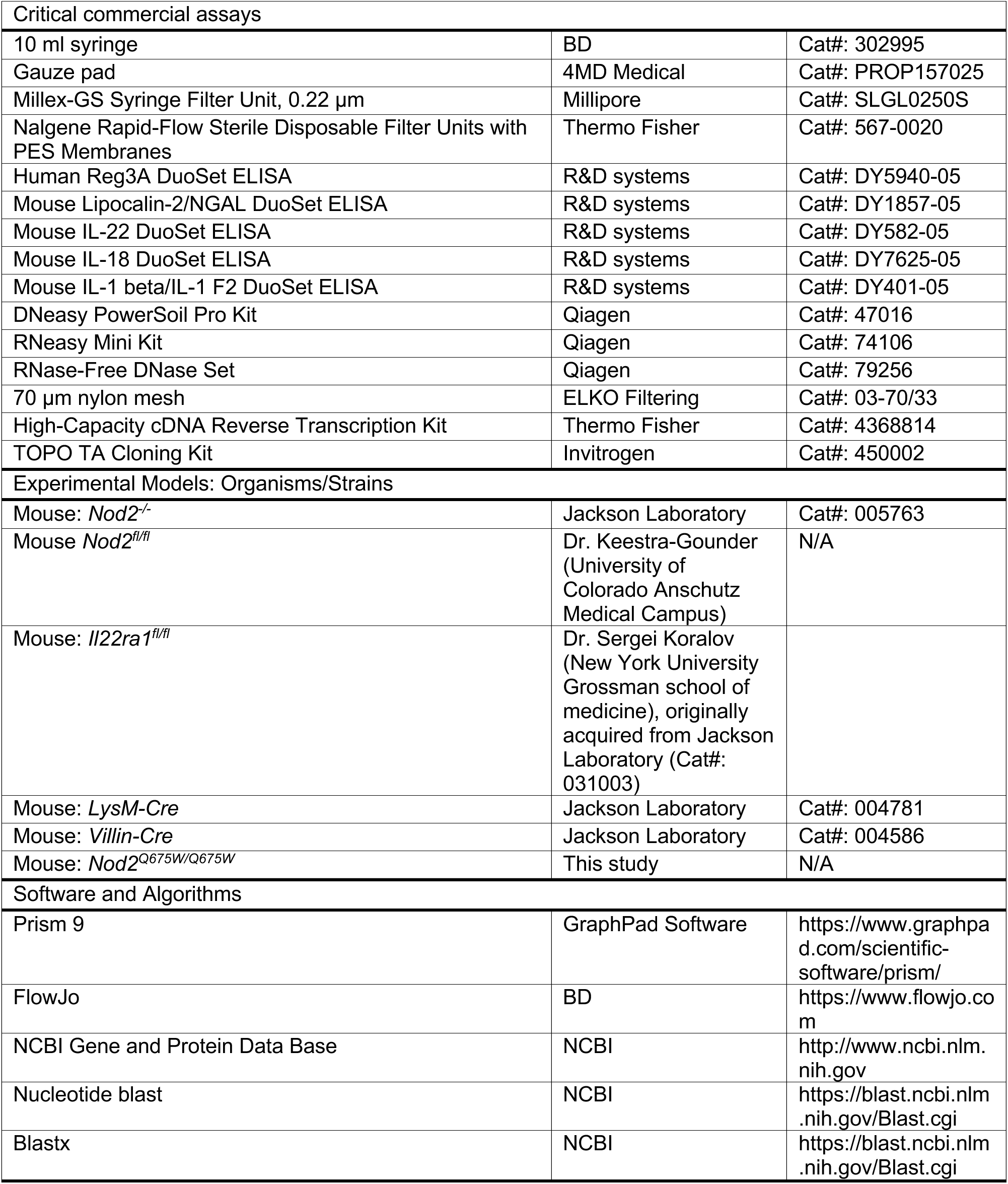
KEY RESOURCES TABLE

## RESOURCE AVAILABILITY

### Lead contact

Further information and requests for resources and reagents should be directed to and will be fulfilled by the Lead Contact, Ken Cadwell.

### Materials availability

The materials in the current study are available from the Lead Contact with a completed Materials Transfer Agreement.

### Data and code availability

The 16S sequencing data are deposited in the NCBI Sequence Read Archive under BioProject accession number PRJNA915760.

## EXPERIENTAL MODEL AND SUBJECT DETAILS

### Mice

Age- and gender-matched 7 to 10-week-old mice on the C57BL/6J (B6) background were used. All mice were bred on-site. Mice harboring gene deletions were compared to their respective littermate control mice indicated in the results section. *Nod2*^-/-^, *LysM-Cre*, and *Villin-Cre* mice were purchased from The Jackson Laboratory. *Il22ra1^fl/fl^* and *Nod2^fl/fl^* mice were obtained from Sergei Koralov (New York University Grossman School of Medicine) and A. Marijke Keestra-Gounder (University of Colorado Anschutz School of Medicine)^78^, respectively. To generate cell type-specific knock-out mice, *Nod2^fl/fl^* mice were bred with *LysM-Cre* and *Villlin-Cre* mice and *Il22ra1^fl/fl^* mice were bred with *Villin-Cre* mice. Mice were assigned numbers to facilitate blind data collection and distributed randomly to treatment groups. All animal studies were performed according to approved protocols by the NYU Grossman School of Medicine Animal Care and Use Committee (IACUCs).

### Generation of *Nod2 Q675W* knock-in mice

CRISPR–Cas9 gene-targeting mixture containing sgRNA (5’-AGCGGGCACGTGCCTGACGC-3’) targeting exon 4 of B6 *Nod2* and template (5’-TCACAGCAGCCTTCCTAGCAGGTCTGTTGTCCCAGCAGCATCGGGACCTGTTGGCTGCATGCCAGGTCTCCGAGAGGGTACTTCTATGGCGTCAGGCACGTGCCCGCTCGTGTCTGGCCCACAGCCT-3’) (synthesized at Integrated DNA Technologies) and Cas9 mRNA were injected into the cytoplasm of zygotes generated from B6 females impregnated by B6 males, and then the microinjected embryos were incubated in potassium-supplemented simplex optimized medium (KSOM) at 37°C for one day and subsequently transferred into pseudopregnant CD-1 female mice at the two-cell stage by the Rodent Genetic Engineering Laboratory at NYU Grossman School of Medicine. The resulting F0 chimeras were screened through genotyping PCR. Amplicons were generated using a pair of primers (Fwd 5’-CTTTTCAGCTGTGGCCGGCT-3’ and Rev 5’-TTTGCCACAGGCCCAATCGG-3’) flanking the targeting sites from tail DNA from chimeras and wild-type mice and cut by BsaHI (NEB) (Supplemental figure 7A). A 3042-bp region containing the *Nod2* coding region was sequenced to verify correct gene targeting (Supplemental figure 7C). All mice used in experiments were backcrossed with B6 mice at least 3 generations.

### Human stool samples and data collection

Stool samples were collected with consent from hospitalized adult non-IBD (NIBD) and IBD patients with gastrointestinal symptoms (Supplemental tables 1 and 2) with consent. The protocol has been approved by the New York University School of Medicine Institutional Review Board (Mucosal Immune Profiling in Patients with Inflammatory Bowel Disease; S12-01137). The clinical data of NIBD and IBD patients were collected using EPIC EHR and REDCap 9.3.6 software. At the time of sample acquisition and processing, investigators were blinded to the patient clinical status.

## METHOD DETAILS

### Bacterial species

*Enterococcus faecalis* OG1RF (*Efl*)*, E. faecium* Com15 (*Efm*), and *Lactococcus lactis thyA* auxotroph (*Lls*) expressing wild-type SagA (*Lls*-*SagA^WT^*) and catalytic mutant SagA (*Lls*-*SagA^C443A^*) were previously described^50^. *Salmonella* Typhimurium SL1344 (*S*Tm) was provided by Dan Littman (NYU). *Efl* and *Efm* were grown at 37°C under ambient atmosphere in autoclaved antibiotics-free Bacto Brain Heart Infusion (BHI) broth (BD). *Lls* strains and *S*Tm were cultured at the above condition in M17 broth (BD) supplemented with 2% (v/v) lactose (Sigma-Aldrich) and 20 μg/ml of thymidine (Sigma-Aldrich) (LM17 broth) and Luria Bertani (LB) broth (NYU Reagent Preparation Core), respectively. Colony forming unit (CFU) of *Efl*, *Efm*, *Lls* strains, and *S*Tm was enumerated on bile esculin azide (BEA) agar (Millipore), HiCrome *Enterococcus faecium* Agar with Selective Supplement (HIMEDIA), M17 agar (BD) supplemented with 2% (v/v) lactose and 20 μg/ml of thymidine (LM17 agar), and LB agar (NYU Reagent Preparation Core), respectively.

### Preparation of human stool extract, antimicrobial activity assay, and REG3A measurement

1 g of human stools from NIBD and IBD patients were homogenized in 5 ml of phosphate-buffered saline (PBS, Corning) using TissueRuptor (Qiagen). 0.5-1 ml of human stool slurries were taken for quantification of bacterial burden. The remaining stool slurries were filtered using a 10 ml syringe (BD) with gauze (4MD Medical). Human stool extracts were collected by centrifugation at 15,000 x *g* for 20 min at 4°C and filter-sterilized using a 10 ml syringe with Millex-GS Syringe Filter Unit, 0.22 μm (Millipore).

For quantifying the antimicrobial activity of the human stool extracts, overnight cultures of *Efl*, *Efm*, and *S*Tm were harvested, resuspended in 50 μl of PBS, and mixed with one volume of human stool extracts or PBS. The mixture of bacteria and human stool extract was incubated in a 37°C static incubator for 24 h and then plated in serial dilution on selective agars as mentioned above for enumerating *Efl*, *Efm*, and *S*Tm.

Quantification of REG3A in human stool extracts was performed using Human Reg3A DuoSet ELISA (R&D systems) according to the manufacturer’s instructions.

### Western blotting

Mouse colonic tissues were processed for immunoblotting as previously described^79, 80^. Briefly, proximal colonic tissues (2 mm) were cut open and washed with PBS, then suspended in lysis buffer (20 mM Tris-HCl (pH 7.4, NYU Reagent Preparation Core), 150 mM NaCl (NYU Reagent Preparation Core), 1% Triton X-100 (Sigma-Aldrich), 10% glycerol (Sigma-Aldrich), and 2x Halt Protease and Phosphatase Inhibitor Cocktail (Thermo Fisher)) and homogenized using FastPrep-24 Classic bead beating grinder and lysis system (MP Biomedicals). Tissue homogenate was then pelleted twice at 10,000 x *g* for 10 min at 4°C to collect the lysates. human stool extracts and mouse colonic tissue lysates were resolved on Bolt 4-12% Bis-Tris Plus Gels (Invitrogen), transferred onto polyvinylidene difluoride membranes, and blocked using Intercept (TBS) blocking buffer (LI-COR). Membranes were probed with primary antibody overnight at 4°C. The following primary antibodies were used for western blotting studies: anti-REG1A (R&D systems, MAB4937), REG3A (R&D systems, MAB5965), REG3G (Abcam, ab233480), REG4 (Abcam, ab255820), NOD2 (Invitrogen, MA1-16611), and β-actin (Sigma-Aldrich, A5441). After incubation with the primary antibody, the membrane was washed and probed with the secondary antibody for 1 h at room temperature (RT). As for secondary antibodies, IRDye 680RD Goat anti-Rabbit (925-68071), IRDye 800CW Goat anti-Mouse (925-32210), and IRDye 800CW Goat anti-Rat (925-32219) were purchased from LI-COR. After additional washing, the protein was then detected with Image Studio for Oddyseey CLx (LI-COR).

### DNA extraction and 16S rRNA sequencing analysis

DNA from human stool samples was extracted with DNeasy PowerSoil Pro kit (Qiagen) according to the manufacturer’s instruction. Bacterial 16S rRNA gene was amplified at the V4 region using primer pairs and paired-end amplicon sequencing was performed on the Illumina MiSeq system at NYU Genome Technology Core. Sequencing reads were processed using the DADA2 pipeline in the QIIME2 software package. Taxonomic assignment was performed against the Greengenes 13_8 99% OTUs full-length sequences database^81^. Alpha diversity analysis was done using observed OTUs^82^. Beta diversity was calculated using Bay-Curtis, Jaccard, unweighted UniFrac, and weighted UniFrac distance and visualized with EMPeror^83^.

### Enumeration of total *Enterococcus* and *Efm* in human stool samples

The human stool slurries (200 mg/ml) were plated in serial dilution on selective agars mentioned above for enumerating CFU of total *Enterococcus* and *Efm*, respectively.

### Bacterial inoculation and DSS treatment of mice

*Efm* drinking water was prepared as described previously^59^. Briefly, overnight culture of *Efm* was harvested and resuspended in filter-sterilized drinking water to 10^9^ CFU/ml, herein referred as *Efm* water, and administered to mice for 14 days. This drinking water was then replaced with one containing 5% dextran sulfate sodium (DSS, TdB Consultancy) for 6 days and then switched to regular drinking water for the remainder of the experiment when examining intestinal injury following *Efm* colonization. For experiments in which *Efm* was administered after the initiation of intestinal injury, mice were treated with 5% DSS for 6 days and received *Efm* water or control water for the remainder of the experiment. Drinking water was exchanged with freshly prepared ones every 2-3 days for all types of treatment.

Overnight cultures of *Lls* strains were harvested and resuspended in PBS to 10^9^ CFU/100 μl. Water containing ampicillin (1 mg/ml, American Bioanalytical) and streptomycin (0.5 mg/ml, Sigma-Aldrich) was filter-sterilized using Nalgene Rapid-Flow Sterile Disposable Filter Units with PES Membranes (Thermo Fisher) for the antibiotic (abx) treatment. The mice treated with abx-containing water for 7 days were given 3% DSS for 6 days. On days 8 and 14, the mice were orally administered 100 μl inoculum of *Lls* strains (approximately 10^9^ CFU).

For treatment of *Lls* culture supernatant, overnight cultures of *Lls* strains were centrifuged at 6,000 x *g* for 15 min at 4°C to collect the culture supernatant. The supernatants and LM17 broth control were filter-sterilized using Nalgene Rapid-Flow Sterile Disposable Filter Units with PES Membranes to avoid bacterial contamination. The mice receiving the filter-sterilized supernatants of *Lls* strains or LM17 broth as drinking water for 7 days were given 5% DSS for 6 days.

The mice were monitored daily for survival and weight loss. Disease score was quantified based on five parameters in which eight was the maximum score: diarrhea (0-2), bloody stool (0-1), hunched posture (0-2), mobility (0-2), and fur ruffling (0-1).

### Quantification of bacterial burden and lipocalin-2 (LCN2) in murine stool samples

Stool pellets from individual mice were weighed, homogenized in PBS, and plated in serial dilution on selective agars as mentioned above for enumerating CFU of total *Enterococcus*, *Efm*, and *Lls* strains, respectively. The remaining stool homogenates were centrifuged at 15,000 x *g* for 5 min to collect clear supernatants. LCN2 in the clear supernatants was quantified using Mouse Lipocalin-2/NGAL DuoSet ELISA (R&D systems) according to the manufacturer’s instructions.

### Histology

Quantification of colon histology data was performed blind. Colonic tissues were cut open along the length, pinned on black wax, and fixed in 10% formalin (Thermo Fisher). Tissues were embedded in 3% low melting point agar (Promega). Formalin embedding, cutting, and hematoxylin and eosin (H&E) staining were performed by the NYU Histopathology core. Sections were imaged on a Nikon Eclipse Ci microscope. Pathologic changes in colonic mucosa evaluated by Y.D. included colonic epithelium, lamina propria, muscularis mucosa, and submucosa. It was based on the state of the changes (acute or chronic) and the degree of involvement. Acute changes primarily included neutrophils in the crypt lumen, erosions, ulcers, pus, and the formation of polyps. Features of chronic changes included the following: crypt distortion (sideways crypts, branching, tortuous), crypt loss, crypt atrophy, basal plasmacytosis, and lymphoid aggregates. The involvement pattern included focal distal involvement, patchy involvement with skipping areas, diffuse involvement, and pancolonic (no skip areas) involvement. The percentage of involvement was calculated as 100 x (Length of disease involved colon / Total length of the colon).

### Colon explant culture and cytokine detection

Colon explant culture was performed as previously described^66^. The proximal colon tissue (1 cm) was opened longitudinally, washed with PBS and cultured for 48 h in 1ml of complete Roswell Park Memorial Institute (RPMI) 1640 (Gibco) containing 2 mM L-glutamine (Corning), penicillin/streptomycin solution (Corning), 20 mM N-2-hydroxyethylpiperazine-N’-2-ethane sulfonic acid (HEPES, Corning), Minimum Essential Medium (MEM) non-essential amino acids (Corning), 1 mM sodium pyruvate (Corning), and β-mercaptoethanol (Gibco). Cytokines in supernatants were measured using the Mouse IL-22, IL-18, and IL-1 beta/IL-1 F2 DuoSet ELISA (R&D systems) according to the manufacturer’s instructions.

### Isolation of lamina propria (LP) cells

LP cells were harvested as before^84^. Briefly, colonic tissue including caecum was cut open and washed with PBS, and fat was removed. The tissues were incubated first with 20 ml of Hank’s Balanced Salt Solution (HBSS, Gibco) with 2% HEPES, 1% sodium pyruvate, 5 mM Ethylenediaminetetraacetic acid (EDTA, NYU Reagent Preparation Core), and 1mM DL-dithiothreitol (Sigma-Aldrich) for 15 min at 37°C with 220 rpm and then with new 20 mL of HBSS with 2% HEPES, 1% sodium pyruvate, 5mM EDTA for 10 min at 37°C with 210 rpm. The tissue bits were washed with HBSS, minced, and then enzymatically digested with collagenase (0.5 mg/ml, Sigma-Aldrich) and Deoxyribonuclease I (0.01 mg/ml, Sigma-Aldrich) for 30 min at 37°C with 200 rpm. Digested solutions were passed through 70 μm nylon mesh (ELKO Filtering) and isolated cells were resuspended in 40% Percoll (Sigma-Aldrich), layered onto 80% Percoll, and centrifuged at RT at 2,200 rpm for 22 min. Cells were recovered from the interphase and washed with RPMI containing 10% fetal bovine serum (FBS, Peak Serum) (cRPMI). For the analysis of the cytokine production, LPLs were plated in cRPMI and stimulated with 1X Cell Stimulation Cocktail (plus transport inhibitors) (eBiosciecne) for 4 h at 37°C.

### Flow cytometry

Surface and intracellular cytokine staining was performed per manufacturer’s instruction in PBS with 0.5% bovine serum albumin (BSA, Thermo Fisher) (BSA/PBS) for 20 min on ice. Two staining panels were prepared as described previously^84^. The following antibodies were used for the first panel: CD11b (101228, 1:200), CD11c (117328, 1:200), CD127 (135041, 1:33), CD4 (100414, 1:150), CD44 (103030, 1:100), CD62L (104433, 1:100), CD8 (100730, 1:100), GR1 (108428, 1:300), NK1.1 (108736, 1:150), Tbet (644810, 1:50), TCRβ (109243, 1:100), TER119 (116228, 1:200) from Biolegend, CD19 (45-0193-82, 1:100) and FOXP3 (17-5773-82, 1:100) from eBioscience, and CD45 (564279, 1:200), GATA3 (560163, 1:40), RORγt (562894, 1:100), TCRγδ (563532, 1:100) from BD Bioscience. The following antibodies were used for the second panel: CD11b (101228, 1:200), CD11c (117328, 1:200), CD127 (135041, 1:33), CD4 (100414, 1:150), CD8 (100730, 1:100), GR1 (108428, 1:300), NK1.1 (108728, 1:200), TCRβ (109243, 1:100), TER119 (116228, 1:200) from Biolegend, IL-22 (12-7221-82, 1:100) from eBioscience, and CD45 (564279, 1:200), IL-10 (566295, 1:100) and TCRγδ (563532, 1:100) from BD Bioscience.

Samples were fixed with either Fixation Buffer (Biolegend) or Foxp3/Transcription Factor Staining Buffer Set (Thermo Fisher). For intracellular staining of the transcription factor, cells were permeabilized with the Foxp3/Transcription Factor Staining Buffer Set at RT for 45 min in the presence of antibodies. For intracellular staining of cytokines, cells were permeabilized with Intracellular Staining Permeabilization Wash Buffer (Biolegend) at RT for 20 min in the presence of antibodies. Zombie UV Fixable Viability Kit (Biolegend) was used to exclude dead cells. Samples were acquired on a BD LSR II (BD Biosciences) and analyzed using FlowJo software (BD).

### *Nod2* sequencing

*Nod2* mRNA was sequenced as previously described ^85^. Briefly, total RNA was extracted from proximal colonic tissues (2 mm) using RNeasy Mini Kit with DNase treatment (QIAGEN), and synthesis of cDNA was conducted with High-Capacity cDNA Reverse Transcription Kit (Thermo Fisher) according to the manufacturer’s protocol. A 3042-bp region containing the *Nod2* coding region was amplified from cDNA by PCR using a pair of primers (Fwd 5’-ATGTGCTCACAGGAAGAGTTC C-3’ and Rev 5’-TCACAACAAGAGTCTGGCGTCCC-3’).

Amplicons were cloned into pCR2.1-TOPO (Invitrogen). The plasmids were sequenced by Oxford Nanopore from SNPsaurus.

### Statistical analyses

Statistical differences were determined as described in the figure legend using GraphPad Prism 9 software. *p* < 0.05 was considered statistically significant for all assays, and individual *p* values are indicated in the figure panels.

## ACKNOWLEDGEMENT

We would like to acknowledge NYU Grossman School of Medicine Flow Cytometry and Cell Sorting, Microscopy, Genome Technology, Rodent Genetic Engineering, and Histology Cores and Clinical Microbiology Laboratory for use of their instruments and technical assistance (supported by National Institutes of Health [NIH] grants P31CA016087, S10OD01058, and S10OD018338). This work was supported in part by NIH grants DK093668 (K.C.), AI121244 (K.C.), HL123340 (K.C.), AI130945 (K.C.), AI140754 (K.C.), DK124336 (K.C.), AI164154 (A.M.K-G.), AI173121 (A.M.K-G.), DK122698 (F.Y.), and DK132908 (M.L.); Faculty Scholar grant from the Howard Hughes Medical Institute (K.C.), Crohn’s & Colitis Foundation (K.C.), Kenneth Rainin Foundation (H.C.H. and K.C.), Bernard Levine Postdoctoral Research Fellowship in Immunology (Y.H.C.) and the Charles H. Revson Senior Fellowships in Biomedical Science (Y.H.C.).

## COMPETING INTERESTS

K.C. has received research support from Pfizer, Takeda, Pacific Biosciences, Genentech, and Abbvie. K.C. has consulted for or received an honoraria from Puretech Health, Genentech, and Abbvie. K.C. is an inventor on U.S. patent 10,722,600 and provisional patent 62/935,035 and 63/157,225. H.C.H. has received research support from Rise Therapeutics and LISCure Biosciences. U.S. patents PCT/US16/28836 (H.C.H.) and PCT/US2020/019038 (H.C.H. and M.E.G.) were obtained for the commercial use of SagA-engineered bacteria. Rise Therapeutics has licensed these patents to develop SagA-probiotics as therapeutics. J.A. reports consultancy fees, honoraria, or advisory board fees from Abbvie, Adiso, Bristol Myers Squibb, Janssen, Pfizer, Fresnius, and BioFire Diagnostics.

## AUTHOR CONTRIBUTIONS

K.K.J. and K.C. conceived the study and designed the experiments. K.K.J. contributed to all the experiments and data analysis. T.H., M.L., and Y.C. helped perform mouse experiments. Y.D. performed histopathology analysis. F.Y. generated *Nod2 Q675W* mice. D.E. performed 16S rRNA sequencing analysis. J.A. and S.G. assisted with the design, analysis, and interpretation of clinical data. M.E.G. and H.C.H. provided *Lls* and *Efm* strains and associated technical assistance. A.M.K-G. provided *NOD2^fl/fl^* mice. K.C. oversaw the analysis and interpretation of all experiments. K.K.J., Y. D., and K.C. wrote the manuscript, and all authors commented on the manuscript, data, and conclusion.

**Table S1.** Supplemental Table 1. **Information for IBD patients.** M; male, F; female; CD; Crohn’s disease; UC; ulcerative colitis; CDAI, Crohn’s disease activity index; N/A, not available.

| **Name** | **Sex** | **Age** | **Disease** | **CDAI** | **Total Mayo** |
| --- | --- | --- | --- | --- | --- |
| IBD1 | F | 24 | CD | 250 |  |
| IBD2 | F | 28 | CD | 192 |  |
| IBD3 | M | 49 | CD | 234 |  |
| IBD4 | M | 33 | CD | 258 |  |
| IBD5 | F | 49 | CD | 180 |  |
| IBD6 | M | 26 | CD | 265 |  |
| IBD7 | F | 43 | CD | 260 |  |
| IBD8 | F | 60 | CD | 176 |  |
| IBD9 | F | 36 | CD | 310 |  |
| IBD10 | F | 56 | CD | 323 |  |
| IBD11 | M | 36 | CD | 268 |  |
| IBD12 | M | 21 | CD | 397 |  |
| IBD13 | M | 19 | CD | 261 |  |
| IBD14 | M | 36 | CD | 251 |  |
| IBD15 | F | 35 | CD | 187 |  |
| IBD16 | F | 78 | CD | 235 |  |
| IBD17 | F | 36 | CD | 275 |  |
| IBD18 | F | 62 | CD | 313 |  |
| IBD19 | F | 20 | CD | 104 |  |
| IBD20 | M | 28 | CD | 298 |  |
| IBD21 | F | 58 | CD | 275 |  |
| IBD22 | F | 33 | CD | 300 |  |
| IBD23 | F | 51 | CD | 372 |  |
| IBD24 | M | 25 | CD | 292 |  |
| IBD25 | M | 42 | CD | 236 |  |
| IBD26 | M | 36 | CD | 301 |  |
| IBD27 | F | 56 | CD | 358 |  |
| IBD28 | M | 30 | CD | 87 |  |
| IBD29 | F | 58 | CD | 264 |  |
| IBD30 | M | 28 | CD | 337 |  |
| IBD31 | M | 69 | CD | 341 |  |
| IBD32 | M | 30 | UC |  | 10 |
| IBD33 | M | 41 | UC |  | 7 |
| IBD34 | F | 58 | UC |  | 8 |
| IBD35 | M | 40 | UC |  | 10 |
| IBD36 | M | 42 | UC |  | 9 |
| IBD37 | M | 30 | UC |  | 10 |
| IBD38 | M | 65 | UC |  | 10 |
| IBD39 | M | 34 | UC |  | 10 |
| IBD40 | M | 28 | UC |  | 7 |
| IBD41 | M | 34 | UC |  | 10 |
| IBD42 | M | 78 | UC |  | 11 |
| IBD43 | M | 33 | UC |  | 10 |
| IBD44 | F | 28 | UC |  | 11 |
| IBD45 | F | 26 | UC |  | 9 |
| IBD46 | F | 35 | UC |  | 11 |
| IBD47 | M | 62 | UC |  | 4 |
| IBD48 | F | 27 | UC |  | 11 |
| IBD49 | F | 28 | UC |  | 10 |
| IBD50 | F | 36 | UC |  | 10 |
| IBD51 | F | 58 | UC |  | 3 |
| IBD52 | F | 35 | UC |  | 11 |
| IBD53 | M | 41 | UC |  | 9 |
| IBD54 | F | 48 | UC |  | 9 |
| IBD55 | F | 18 | UC |  | 7 |
| IBD56 | M | 66 | UC |  | N/A |

**Table S2.** Supplemental Table 2. Information for non-IBD (NIBD) patients.

| Name | Sex | Age | GI-related medical history | GI symptoms |
| --- | --- | --- | --- | --- |
| NIBD1 | F | 83 |  | Diarrhea for 7 days |
| NIBD2 | M | 61 | *Clostridia difficile* infection | Intermittent diarrhea for 3 months. |
| NIBD3 | M | 37 | Irritable bowel syndrome | Intermittent diarrhea with urgency, bloating and chest discomfort every 2 weeks |
| NIBD4 | F | 50 | Hirschsprung's disease, type 2 diabetes mellitus | Vomiting and watery diarrhea and for 4 days |
| NIBD5 | F | 40 |  | Nausea, Vomiting, and diarrhea for 5 days |
| NIBD6 | M | 86 | Whipple procedure | Diarrhea and abdominal pain for 3 days |
| NIBD7 | M | 56 |  | Diarrhea and abdominal pain |
| NIBD8 | F | 42 |  | Bloody diarrhea and abdominal pain |
| NIBD9 | M | 9 |  | Diarrhea |
| NIBD10 | M | 35 |  | Right Lower Quadrant pain, Nausea, Vomiting, and diarrhea for 1 day |
| NIBD11 | F | 29 |  | Abdominal pain |
| NIBD12 | F | 58 |  | Vomiting and abdominal tenderness |
| NIBD13 | F | 44 |  | Nausea, vomiting, and diarrhea for 4 days |
| NIBD14 | M | 23 |  | Nausea, vomiting, and diarrhea with fever for 2 days |
| NIBD15 | M | 75 |  | Diarrhea for 2 weeks |
| NIBD16 | M | 51 |  | Diarrhea for 10 days |
| NIBD17 | M | 70 |  | Diarrhea |
| NIBD18 | F | 49 |  | Diarrhea for 5 days |
| NIBD19 | F | 53 |  | Abdominal pain |
| NIBD20 | F | 54 |  | Constipation and rectal pain |
| NIBD21 | M | 79 |  | Bloody stool |
| NIBD22 | F | 69 |  | Diarrhea |
| NIBD23 | M | 71 |  | Diarrhea |
| NIBD24 | F | 65 | *C. difficile* infection | Nausea, vomiting, and diarrhea for 4 days |
| NIBD25 | F | 53 |  | Nausea and diarrhea for 4 days |
| NIBD26 | F | 49 |  | Diarrhea for 2 weeks |
| NIBD27 | F | 23 | Recurrent *C. difficile* infection | Diarrhea and abdominal pain for 1 week |
| NIBD28 | M | 86 |  | Diarrhea and abdominal pain |
| NIBD29 | F | 74 |  | Diarrhea and abdominal pain with fever for 2 days |
| NIBD30 | F | 63 |  | Vomiting, diarrhea, abdominal pain, neutropenic fever |
| NIBD31 | M | 52 | Pancreatic cancer | Unclear |
| NIBD32 | F | 36 |  | Abdominal pain |
| NIBD33 | F | 73 |  | Abdominal pain and bloating |
| NIBD34 | F | 55 |  | Antibiotic associated diarrhea |
| NIBD35 | F | 33 |  | Vomiting and watery diarrhea |
| NIBD36 | M | 29 |  | Rectal bleeding and bloating |
| NIBD37 | M | 55 |  | Diarrhea for 3 days |
| NIBD38 | F | 72 |  | Constipation with rectal bleeding |
| NIBD39 | F | 38 |  | Nausea, diarrhea, and bloating |
| NIBD40 | F | 76 |  | Diarrhea and abdominal pain |
| NIBD41 | F | 82 |  | Diarrhea |
| NIBD42 | F | 69 |  | Nausea, vomiting, and Diarrhea for 5 days |
| NIBD43 | F | 74 | Microscopic colitis | Diarrhea |
| NIBD44 | F | 29 |  | Rectal bleeding and abdominal pain |
| NIBD45 | F | 55 |  | Diarrhea |
| NIBD46 | M | 37 |  | Loose stools |
| NIBD47 | M | 39 |  | Nausea, vomiting, and diarrhea for 4 days |
| NIBD48 | M | 54 |  | Nausea and watery diarrhea for 6 days |
| NIBD49 | F | 45 |  | Diarrhea |
| NIBD50 | F | 91 | Gastro-Esophageal Reflux Disease | Nausea, vomiting, and diarrhea for 1 day |
| NIBD51 | M | 91 |  | Nausea, vomiting, and diarrhea for 3 days |
| NIBD52 | M | 38 | Chronic norovirus infection | Diarrhea and abdominal pain for 1 week |
| NIBD53 | F | 0 |  | Diarrhea with fever |
| NIBD54 | F | 58 |  | Diarrhea and hematuria for 2 days |
| NIBD55 | F | 75 | *C. difficile* infection and diverticulitis | Nausea and diarrhea for 1 day |
| NIBD56 | F | 51 |  | Abdominal pain |

**Table S3.**
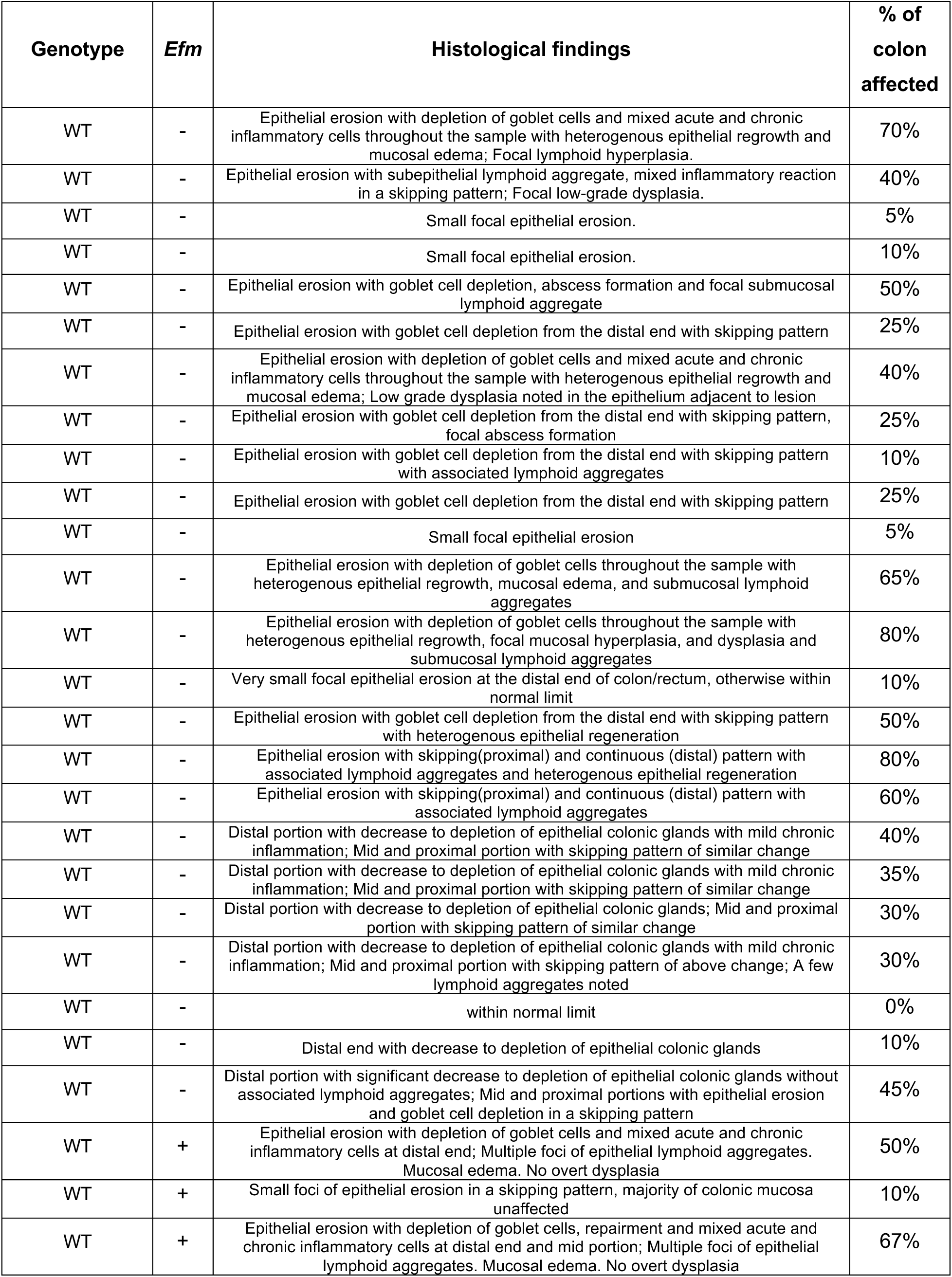
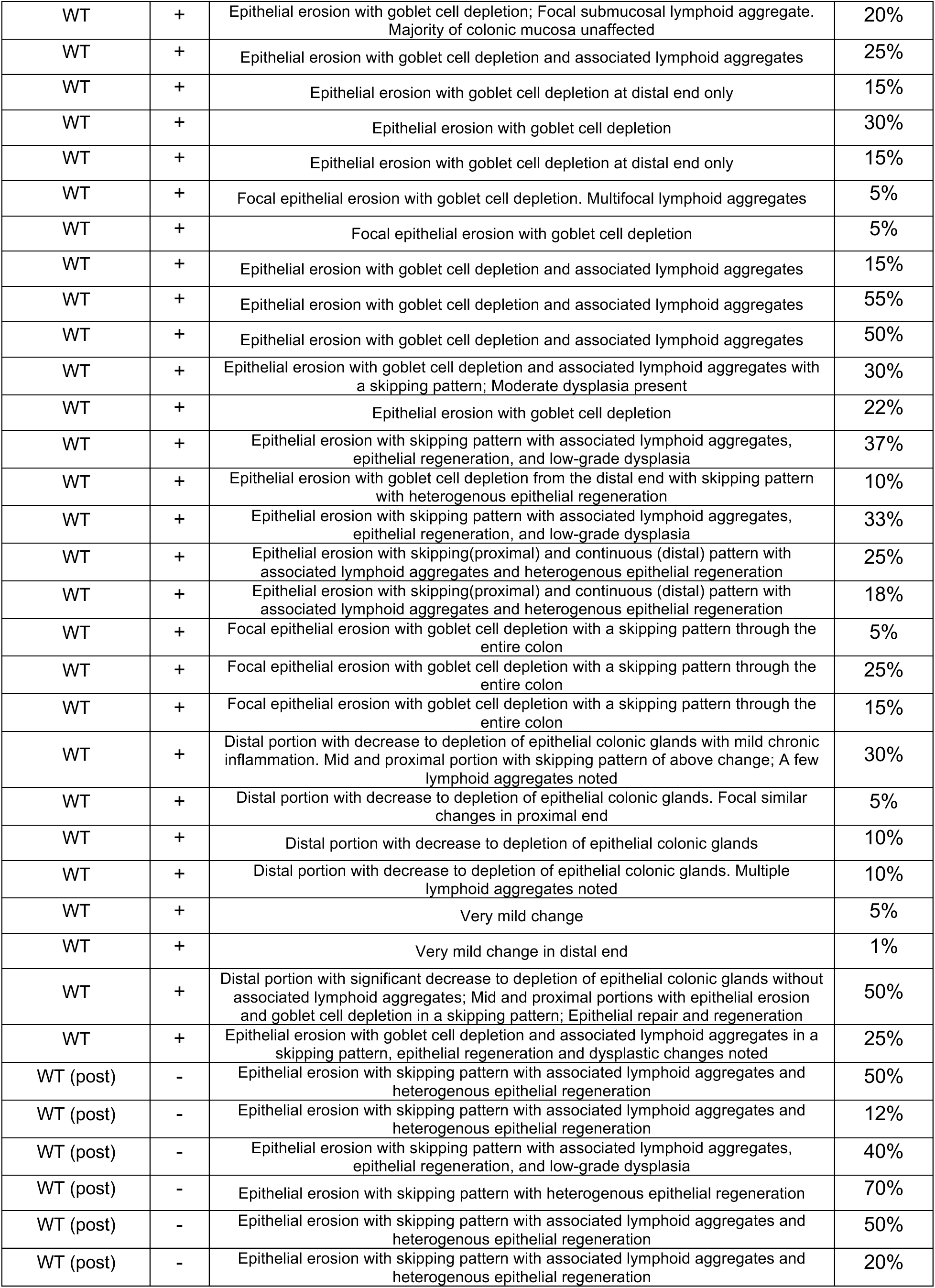
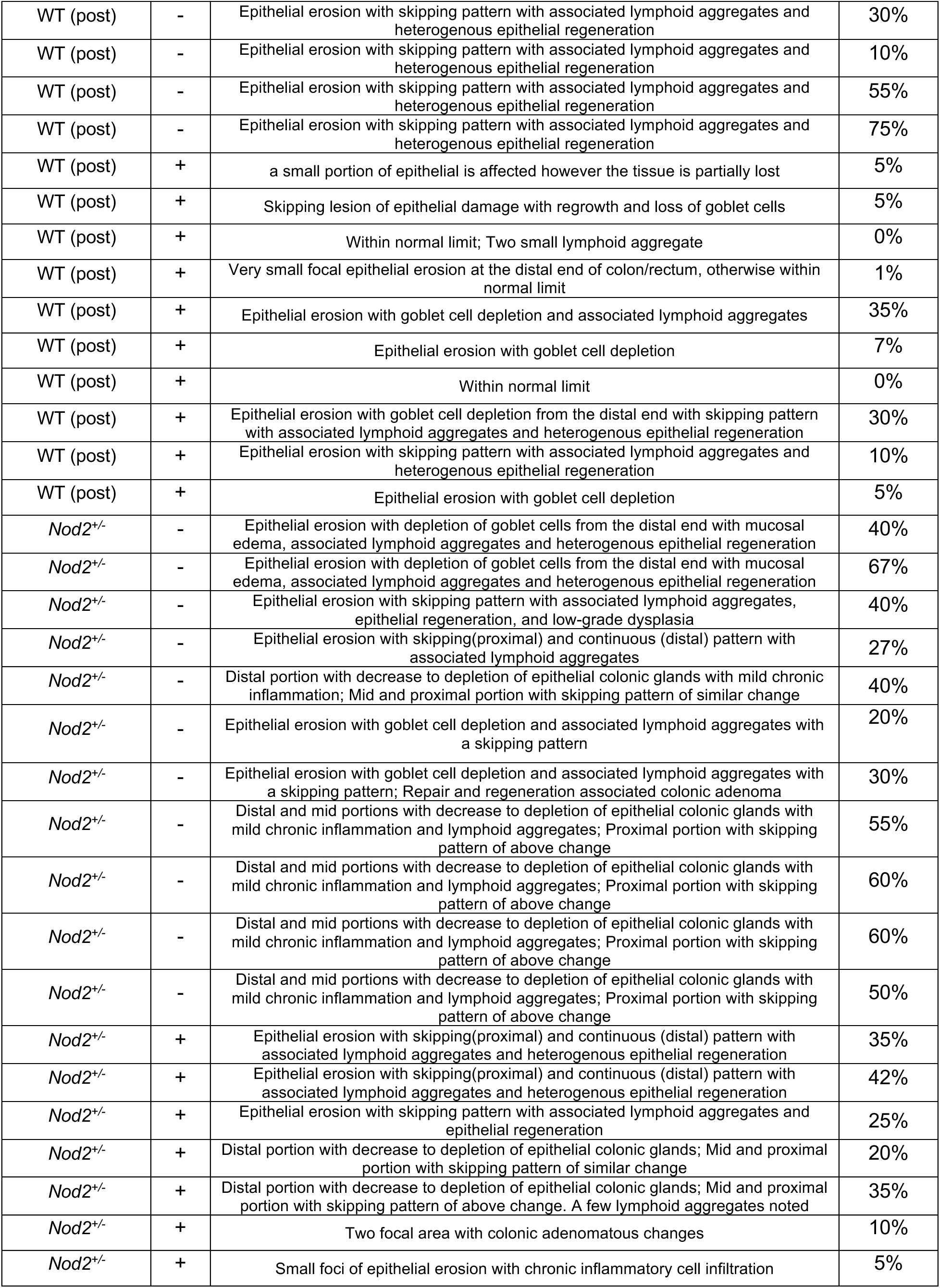
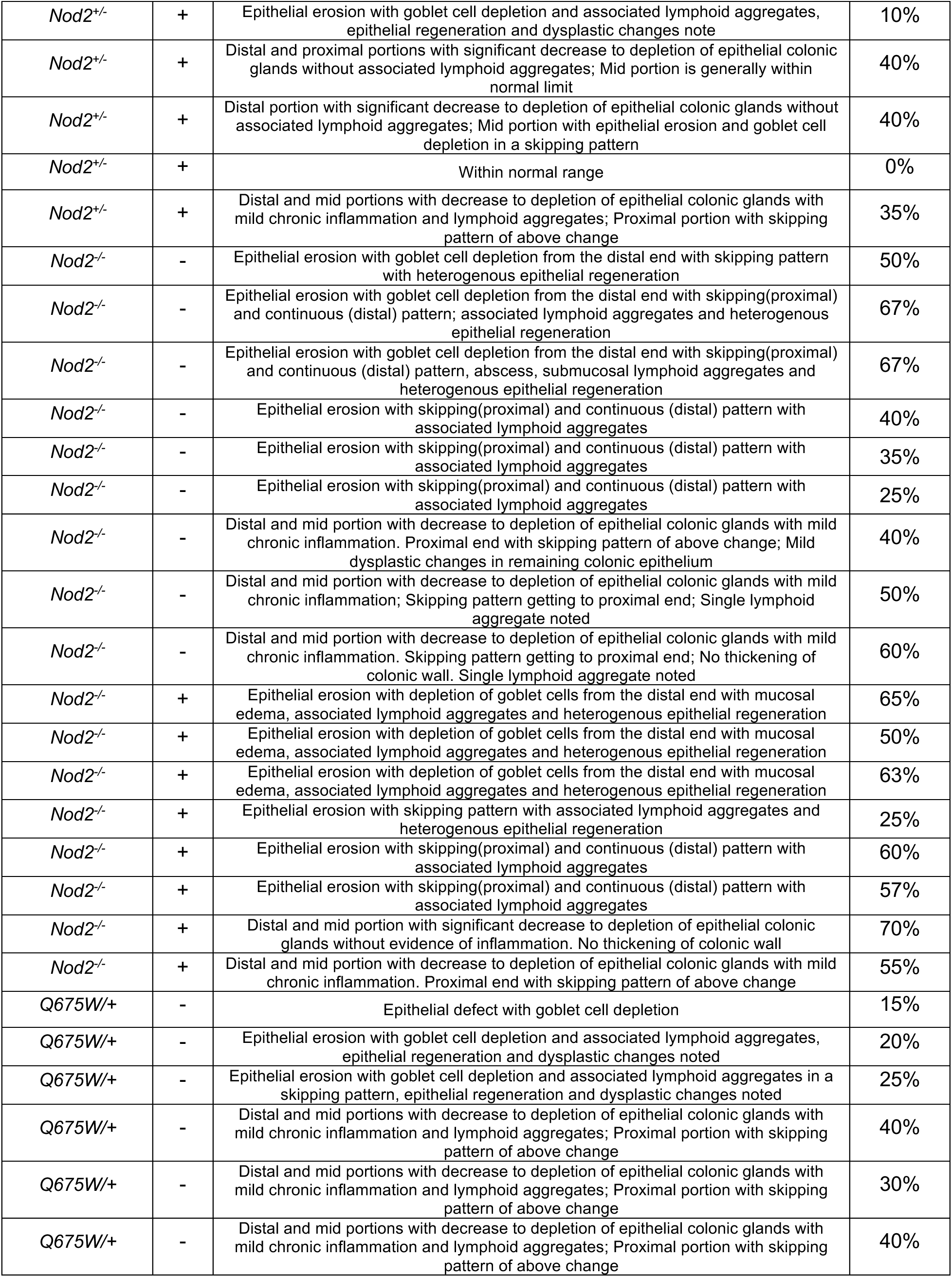
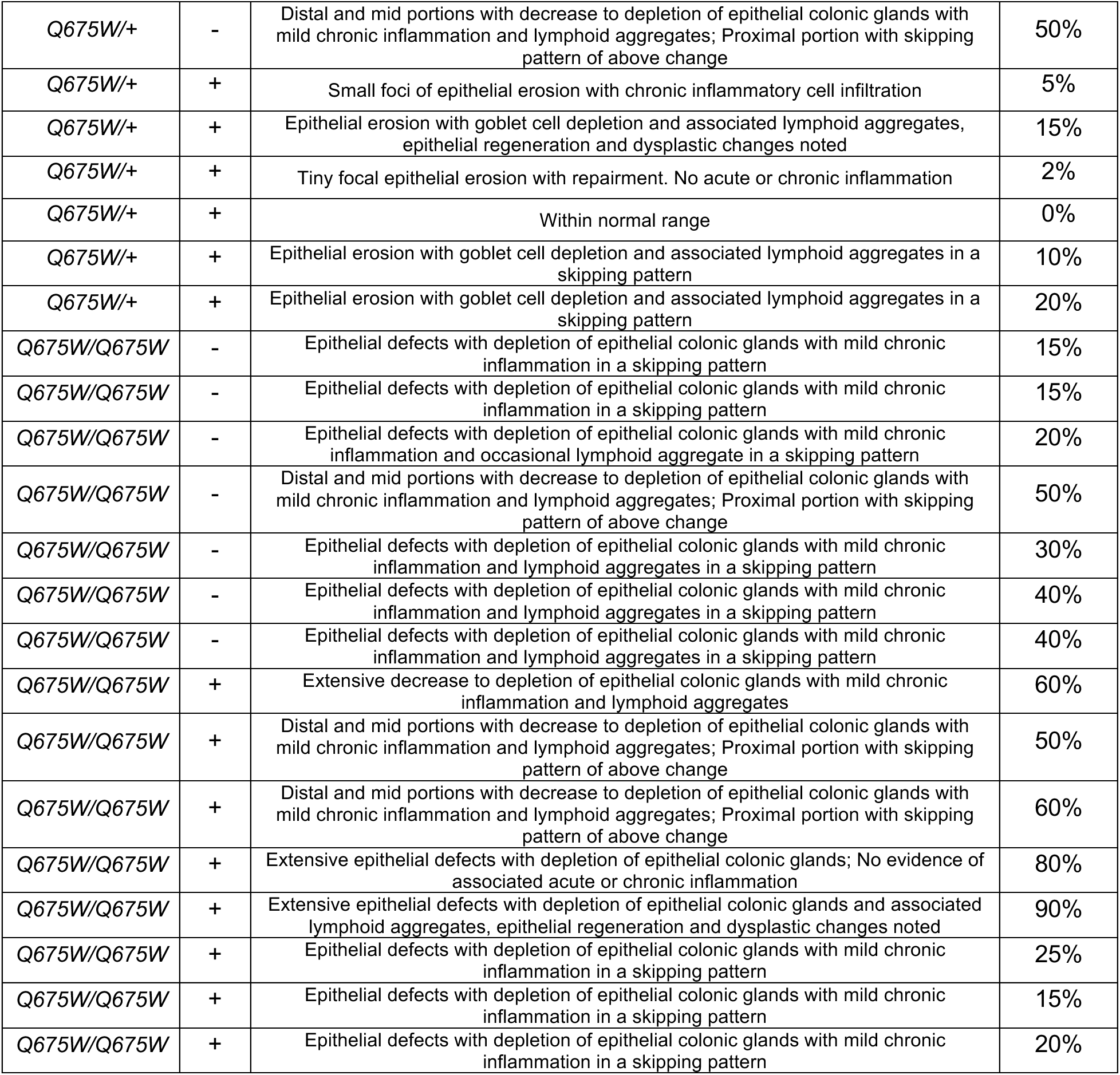
Supplemental Table 3. **Histological findings of colonic sections from mice receiving DSS.** WT, wild type; *Q675W/+*, *Nod2^Q675W/+^*; *Q675W/Q675W*, *Nod2^Q^*^675^*^W/Q675W^*. WT (post) indicated mice receiving *Efm* after DSS treatment.

## Notes

### Summary of Updates

I don't have any revised matters but would like to change distribution/reuse option.

## References

1. Muniz, L.R., Knosp, C., and Yeretssian, G. (2012). Intestinal antimicrobial peptides during homeostasis, infection, and disease. Front Immunol 3, 310. 10.3389/fimmu.2012.00310.

2. Bevins, C.L., and Salzman, N.H. (2011). Paneth cells, antimicrobial peptides and maintenance of intestinal homeostasis. Nat Rev Microbiol 9, 356–368. 10.1038/nrmicro2546.

3. Ramanan, D., and Cadwell, K. (2016). Intrinsic Defense Mechanisms of the Intestinal Epithelium. Cell Host Microbe 19, 434–441. 10.1016/j.chom.2016.03.003.

4. Okumura, R., Kurakawa, T., Nakano, T., Kayama, H., Kinoshita, M., Motooka, D., Gotoh, K., Kimura, T., Kamiyama, N., Kusu, T., et al. (2016). Lypd8 promotes the segregation of flagellated microbiota and colonic epithelia. Nature 532, 117–121. 10.1038/nature17406.

5. Lehotzky, R.E., Partch, C.L., Mukherjee, S., Cash, H.L., Goldman, W.E., Gardner, K.H., and Hooper, L.V. (2010). Molecular basis for peptidoglycan recognition by a bactericidal lectin. Proc Natl Acad Sci U S A 107, 7722–7727. 10.1073/pnas.0909449107.

6. Cash, H.L., Whitham, C.V., Behrendt, C.L., and Hooper, L.V. (2006). Symbiotic bacteria direct expression of an intestinal bactericidal lectin. Science 313, 1126–1130. 10.1126/science.1127119.

7. Wang, L., Fouts, D.E., Starkel, P., Hartmann, P., Chen, P., Llorente, C., DePew, J., Moncera, K., Ho, S.B., Brenner, D.A., et al. (2016). Intestinal REG3 Lectins Protect against Alcoholic Steatohepatitis by Reducing Mucosa-Associated Microbiota and Preventing Bacterial Translocation. Cell Host Microbe 19, 227–239. 10.1016/j.chom.2016.01.003.

8. Vaishnava, S., Yamamoto, M., Severson, K.M., Ruhn, K.A., Yu, X., Koren, O., Ley, R., Wakeland, E.K., and Hooper, L.V. (2011). The antibacterial lectin RegIIIgamma promotes the spatial segregation of microbiota and host in the intestine. Science 334, 255–258. 10.1126/science.1209791.

9. Brandl, K., Plitas, G., Mihu, C.N., Ubeda, C., Jia, T., Fleisher, M., Schnabl, B., DeMatteo, R.P., and Pamer, E.G. (2008). Vancomycin-resistant enterococci exploit antibiotic-induced innate immune deficits. Nature 455, 804–807. 10.1038/nature07250.

10. Darnaud, M., Dos Santos, A., Gonzalez, P., Augui, S., Lacoste, C., Desterke, C., De Hertogh, G., Valentino, E., Braun, E., Zheng, J., et al. (2018). Enteric Delivery of Regenerating Family Member 3 alpha Alters the Intestinal Microbiota and Controls Inflammation in Mice With Colitis. Gastroenterology 154, 1009–1023 e1014. 10.1053/j.gastro.2017.11.003.

11. Zhao, D., Kim, Y.H., Jeong, S., Greenson, J.K., Chaudhry, M.S., Hoepting, M., Anderson, E.R., van den Brink, M.R., Peled, J.U., Gomes, A.L., et al. (2018). Survival signal REG3alpha prevents crypt apoptosis to control acute gastrointestinal graft-versus-host disease. J Clin Invest 128, 4970–4979. 10.1172/JCI99261.

12. Yu, S., Balasubramanian, I., Laubitz, D., Tong, K., Bandyopadhyay, S., Lin, X., Flores, J., Singh, R., Liu, Y., Macazana, C., et al. (2020). Paneth Cell-Derived Lysozyme Defines the Composition of Mucolytic Microbiota and the Inflammatory Tone of the Intestine. Immunity 53, 398–416 e398. 10.1016/j.immuni.2020.07.010.

13. Cadwell, K., Liu, J.Y., Brown, S.L., Miyoshi, H., Loh, J., Lennerz, J.K., Kishi, C., Kc, W., Carrero, J.A., Hunt, S., et al. (2008). A key role for autophagy and the autophagy gene Atg16l1 in mouse and human intestinal Paneth cells. Nature 456, 259–263. 10.1038/nature07416.

14. Wehkamp, J., Salzman, N.H., Porter, E., Nuding, S., Weichenthal, M., Petras, R.E., Shen, B., Schaeffeler, E., Schwab, M., Linzmeier, R., et al. (2005). Reduced Paneth cell alpha-defensins in ileal Crohn’s disease. Proc Natl Acad Sci U S A 102, 18129–18134. 10.1073/pnas.0505256102.

15. Koslowski, M.J., Kubler, I., Chamaillard, M., Schaeffeler, E., Reinisch, W., Wang, G., Beisner, J., Teml, A., Peyrin-Biroulet, L., Winter, S., et al. (2009). Genetic variants of Wnt transcription factor TCF-4 (TCF7L2) putative promoter region are associated with small intestinal Crohn’s disease. PLoS One 4, e4496. 10.1371/journal.pone.0004496.

16. Courth, L.F., Ostaff, M.J., Mailander-Sanchez, D., Malek, N.P., Stange, E.F., and Wehkamp, J. (2015). Crohn’s disease-derived monocytes fail to induce Paneth cell defensins. Proc Natl Acad Sci U S A 112, 14000–14005. 10.1073/pnas.1510084112.

17. Bel, S., Pendse, M., Wang, Y., Li, Y., Ruhn, K.A., Hassell, B., Leal, T., Winter, S.E., Xavier, R.J., and Hooper, L.V. (2017). Paneth cells secrete lysozyme via secretory autophagy during bacterial infection of the intestine. Science 357, 1047–1052. 10.1126/science.aal4677.

18. Manichanh, C., Borruel, N., Casellas, F., and Guarner, F. (2012). The gut microbiota in IBD. Nat Rev Gastroenterol Hepatol 9, 599–608. 10.1038/nrgastro.2012.152.

19. Ni, J., Wu, G.D., Albenberg, L., and Tomov, V.T. (2017). Gut microbiota and IBD: causation or correlation? Nat Rev Gastroenterol Hepatol 14, 573–584. 10.1038/nrgastro.2017.88.

20. Wong, S.Y., and Cadwell, K. (2018). There was collusion: Microbes in inflammatory bowel disease. PLoS Pathog 14, e1007215. 10.1371/journal.ppat.1007215.

21. Jostins, L., Ripke, S., Weersma, R.K., Duerr, R.H., McGovern, D.P., Hui, K.Y., Lee, J.C., Schumm, L.P., Sharma, Y., Anderson, C.A., et al. (2012). Host-microbe interactions have shaped the genetic architecture of inflammatory bowel disease. Nature 491, 119–124. 10.1038/nature11582.

22. Knights, D., Lassen, K.G., and Xavier, R.J. (2013). Advances in inflammatory bowel disease pathogenesis: linking host genetics and the microbiome. Gut 62, 1505–1510. 10.1136/gutjnl-2012-303954.

23. Cho, J.H., and Abraham, C. (2007). Inflammatory bowel disease genetics: Nod2. Annu Rev Med 58, 401–416. 10.1146/annurev.med.58.061705.145024.

24. Bonen, D.K., and Cho, J.H. (2003). The genetics of inflammatory bowel disease. Gastroenterology 124, 521–536. 10.1053/gast.2003.50045.

25. Kim, Y.G., Kamada, N., Shaw, M.H., Warner, N., Chen, G.Y., Franchi, L., and Nunez, G. (2011). The Nod2 sensor promotes intestinal pathogen eradication via the chemokine CCL2-dependent recruitment of inflammatory monocytes. Immunity 34, 769–780. 10.1016/j.immuni.2011.04.013.

26. Zhang, Q., Pan, Y., Yan, R., Zeng, B., Wang, H., Zhang, X., Li, W., Wei, H., and Liu, Z. (2015). Commensal bacteria direct selective cargo sorting to promote symbiosis. Nat Immunol 16, 918–926. 10.1038/ni.3233.

27. Petnicki-Ocwieja, T., Hrncir, T., Liu, Y.J., Biswas, A., Hudcovic, T., Tlaskalova-Hogenova, H., and Kobayashi, K.S. (2009). Nod2 is required for the regulation of commensal microbiota in the intestine. Proc Natl Acad Sci U S A 106, 15813–15818. 10.1073/pnas.0907722106.

28. Ramanan, D., Tang, M.S., Bowcutt, R., Loke, P., and Cadwell, K. (2014). Bacterial sensor Nod2 prevents inflammation of the small intestine by restricting the expansion of the commensal Bacteroides vulgatus. Immunity 41, 311–324. 10.1016/j.immuni.2014.06.015.

29. Marchiando, A.M., Ramanan, D., Ding, Y., Gomez, L.E., Hubbard-Lucey, V.M., Maurer, K., Wang, C., Ziel, J.W., van Rooijen, N., Nunez, G., et al. (2013). A deficiency in the autophagy gene Atg16L1 enhances resistance to enteric bacterial infection. Cell Host Microbe 14, 216–224. 10.1016/j.chom.2013.07.013.

30. Wong, S.Y., Coffre, M., Ramanan, D., Hines, M.J., Gomez, L.E., Peters, L.A., Schadt, E.E., Koralov, S.B., and Cadwell, K. (2018). B Cell Defects Observed in Nod2 Knockout Mice Are a Consequence of a Dock2 Mutation Frequently Found in Inbred Strains. J Immunol 201, 1442–1451. 10.4049/jimmunol.1800014.

31. de Souza, P.R., Guimaraes, F.R., Sales-Campos, H., Bonfa, G., Nardini, V., Chica, J.E.L., Turato, W.M., Silva, J.S., Zamboni, D.S., and Cardoso, C.R.B. (2018). Absence of NOD2 receptor predisposes to intestinal inflammation by a deregulation in the immune response in hosts that are unable to control gut dysbiosis. Immunobiology 223, 577–585. 10.1016/j.imbio.2018.07.003.

32. Robertson, S.J., Geddes, K., Maisonneuve, C., Streutker, C.J., and Philpott, D.J. (2016). Resilience of the intestinal microbiota following pathogenic bacterial infection is independent of innate immunity mediated by NOD1 or NOD2. Microbes Infect 18, 460–471. 10.1016/j.micinf.2016.03.014.

33. Geddes, K., Rubino, S., Streutker, C., Cho, J.H., Magalhaes, J.G., Le Bourhis, L., Selvanantham, T., Girardin, S.E., and Philpott, D.J. (2010). Nod1 and Nod2 regulation of inflammation in the Salmonella colitis model. Infect Immun 78, 5107–5115. 10.1128/IAI.00759-10.

34. Couturier-Maillard, A., Secher, T., Rehman, A., Normand, S., De Arcangelis, A., Haesler, R., Huot, L., Grandjean, T., Bressenot, A., Delanoye-Crespin, A., et al. (2013). NOD2-mediated dysbiosis predisposes mice to transmissible colitis and colorectal cancer. J Clin Invest 123, 700–711. 10.1172/JCI62236.

35. Wehkamp, J., Harder, J., Weichenthal, M., Schwab, M., Schaffeler, E., Schlee, M., Herrlinger, K.R., Stallmach, A., Noack, F., Fritz, P., et al. (2004). NOD2 (CARD15) mutations in Crohn’s disease are associated with diminished mucosal alpha-defensin expression. Gut 53, 1658–1664. 10.1136/gut.2003.032805.

36. Bevins, C.L., Stange, E.F., and Wehkamp, J. (2009). Decreased Paneth cell defensin expression in ileal Crohn’s disease is independent of inflammation, but linked to the NOD2 1007fs genotype. Gut 58, 882–883; discussion 883-884.

37. VanDussen, K.L., Liu, T.C., Li, D., Towfic, F., Modiano, N., Winter, R., Haritunians, T., Taylor, K.D., Dhall, D., Targan, S.R., et al. (2014). Genetic variants synthesize to produce paneth cell phenotypes that define subtypes of Crohn’s disease. Gastroenterology 146, 200–209. 10.1053/j.gastro.2013.09.048.

38. van Beelen Granlund, A., Ostvik, A.E., Brenna, O., Torp, S.H., Gustafsson, B.I., and Sandvik, A.K. (2013). REG gene expression in inflamed and healthy colon mucosa explored by in situ hybridisation. Cell Tissue Res 352, 639–646. 10.1007/s00441-013-1592-z.

39. Ogawa, H., Fukushima, K., Naito, H., Funayama, Y., Unno, M., Takahashi, K., Kitayama, T., Matsuno, S., Ohtani, H., Takasawa, S., et al. (2003). Increased expression of HIP/PAP and regenerating gene III in human inflammatory bowel disease and a murine bacterial reconstitution model. Inflamm Bowel Dis 9, 162–170. 10.1097/00054725-200305000-00003.

40. Gao, J., Zhao, X., Hu, S., Huang, Z., Hu, M., Jin, S., Lu, B., Sun, K., Wang, Z., Fu, J., et al. (2022). Gut microbial DL-endopeptidase alleviates Crohn’s disease via the NOD2 pathway. Cell Host Microbe 30, 1435–1449 e1439. 10.1016/j.chom.2022.08.002.

41. Kim, B., Wang, Y.C., Hespen, C.W., Espinosa, J., Salje, J., Rangan, K.J., Oren, D.A., Kang, J.Y., Pedicord, V.A., and Hang, H.C. (2019). Enterococcus faecium secreted antigen A generates muropeptides to enhance host immunity and limit bacterial pathogenesis. Elife 8. 10.7554/eLife.45343.

42. Kim, Y.G., Shaw, M.H., Warner, N., Park, J.H., Chen, F., Ogura, Y., and Nunez, G. (2011). Cutting edge: Crohn’s disease-associated Nod2 mutation limits production of proinflammatory cytokines to protect the host from Enterococcus faecalis-induced lethality. J Immunol 187, 2849–2852. 10.4049/jimmunol.1001854.

43. Garcia-Solache, M., and Rice, L.B. (2019). The Enterococcus: a Model of Adaptability to Its Environment. Clin Microbiol Rev 32. 10.1128/CMR.00058-18.

44. Balish, E., and Warner, T. (2002). Enterococcus faecalis induces inflammatory bowel disease in interleukin-10 knockout mice. Am J Pathol 160, 2253–2257. 10.1016/S0002-9440(10)61172-8.

45. Seishima, J., Iida, N., Kitamura, K., Yutani, M., Wang, Z., Seki, A., Yamashita, T., Sakai, Y., Honda, M., Yamashita, T., et al. (2019). Gut-derived Enterococcus faecium from ulcerative colitis patients promotes colitis in a genetically susceptible mouse host. Genome Biol 20, 252. 10.1186/s13059-019-1879-9.

46. Ben Braiek, O., and Smaoui, S. (2019). Enterococci: Between Emerging Pathogens and Potential Probiotics. Biomed Res Int 2019, 5938210. 10.1155/2019/5938210.

47. Franz, C.M., Huch, M., Abriouel, H., Holzapfel, W., and Galvez, A. (2011). Enterococci as probiotics and their implications in food safety. Int J Food Microbiol 151, 125–140. 10.1016/j.ijfoodmicro.2011.08.014.

48. Rangan, K.J., Pedicord, V.A., Wang, Y.C., Kim, B., Lu, Y., Shaham, S., Mucida, D., and Hang, H.C. (2016). A secreted bacterial peptidoglycan hydrolase enhances tolerance to enteric pathogens. Science 353, 1434–1437. 10.1126/science.aaf3552.

49. Pedicord, V.A., Lockhart, A.A.K., Rangan, K.J., Craig, J.W., Loschko, J., Rogoz, A., Hang, H.C., and Mucida, D. (2016). Exploiting a host-commensal interaction to promote intestinal barrier function and enteric pathogen tolerance. Sci Immunol 1. 10.1126/sciimmunol.aai7732.

50. Griffin, M.E., Espinosa, J., Becker, J.L., Luo, J.D., Carroll, T.S., Jha, J.K., Fanger, G.R., and Hang, H.C. (2021). Enterococcus peptidoglycan remodeling promotes checkpoint inhibitor cancer immunotherapy. Science 373, 1040–1046. 10.1126/science.abc9113.

51. Wang, Y.C., Westcott, N.P., Griffin, M.E., and Hang, H.C. (2019). Peptidoglycan Metabolite Photoaffinity Reporters Reveal Direct Binding to Intracellular Pattern Recognition Receptors and Arf GTPases. ACS Chem Biol 14, 405–414. 10.1021/acschembio.8b01038.

52. Michot, J.M., Bigenwald, C., Champiat, S., Collins, M., Carbonnel, F., Postel-Vinay, S., Berdelou, A., Varga, A., Bahleda, R., Hollebecque, A., et al. (2016). Immune-related adverse events with immune checkpoint blockade: a comprehensive review. Eur J Cancer 54, 139–148. 10.1016/j.ejca.2015.11.016.

53. Parikh, A., Stephan, A.F., and Tzanakakis, E.S. (2012). Regenerating proteins and their expression, regulation and signaling. Biomol Concepts 3, 57–70. 10.1515/bmc.2011.055.

54. Wang, W., Wang, Y., Lu, Y., Zhu, J., Tian, X., Wu, B., Du, J., Cai, W., and Xiao, Y. (2022). Reg4 protects against Salmonella infection-associated intestinal inflammation via adopting a calcium-dependent lectin-like domain. Int Immunopharmacol 113, 109310. 10.1016/j.intimp.2022.109310.

55. Qi, H., Wei, J., Gao, Y., Yang, Y., Li, Y., Zhu, H., Su, L., Su, X., Zhang, Y., and Yang, R. (2020). Reg4 and complement factor D prevent the overgrowth of E. coli in the mouse gut. Commun Biol 3, 483. 10.1038/s42003-020-01219-2.

56. Chassaing, B., Srinivasan, G., Delgado, M.A., Young, A.N., Gewirtz, A.T., and Vijay-Kumar, M. (2012). Fecal lipocalin 2, a sensitive and broadly dynamic non-invasive biomarker for intestinal inflammation. PLoS One 7, e44328. 10.1371/journal.pone.0044328.

57. Forster, S.C., Clare, S., Beresford-Jones, B.S., Harcourt, K., Notley, G., Stares, M.D., Kumar, N., Soderholm, A.T., Adoum, A., Wong, H., et al. (2022). Identification of gut microbial species linked with disease variability in a widely used mouse model of colitis. Nat Microbiol 7, 590–599. 10.1038/s41564-022-01094-z.

58. Moon, C., Baldridge, M.T., Wallace, M.A., D, C.A., Burnham, Virgin, H.W., and Stappenbeck, T.S. (2015). Vertically transmitted faecal IgA levels determine extra-chromosomal phenotypic variation. Nature 521, 90–93. 10.1038/nature14139.

59. Kommineni, S., Bretl, D.J., Lam, V., Chakraborty, R., Hayward, M., Simpson, P., Cao, Y., Bousounis, P., Kristich, C.J., and Salzman, N.H. (2015). Bacteriocin production augments niche competition by enterococci in the mammalian gastrointestinal tract. Nature 526, 719–722. 10.1038/nature15524.

60. Teng, F., Kawalec, M., Weinstock, G.M., Hryniewicz, W., and Murray, B.E. (2003). An Enterococcus faecium secreted antigen, SagA, exhibits broad-spectrum binding to extracellular matrix proteins and appears essential for E. faecium growth. Infect Immun 71, 5033–5041. 10.1128/IAI.71.9.5033-5041.2003.

61. Rakoff-Nahoum, S., Paglino, J., Eslami-Varzaneh, F., Edberg, S., and Medzhitov, R. (2004). Recognition of commensal microflora by toll-like receptors is required for intestinal homeostasis. Cell 118, 229–241. 10.1016/j.cell.2004.07.002.

62. Kernbauer, E., Ding, Y., and Cadwell, K. (2014). An enteric virus can replace the beneficial function of commensal bacteria. Nature 516, 94–98. 10.1038/nature13960.

63. Ferrand, A., Al Nabhani, Z., Tapias, N.S., Mas, E., Hugot, J.P., and Barreau, F. (2019). NOD2 Expression in Intestinal Epithelial Cells Protects Toward the Development of Inflammation and Associated Carcinogenesis. Cell Mol Gastroenterol Hepatol 7, 357–369. 10.1016/j.jcmgh.2018.10.009.

64. Mukherjee, T., Hovingh, E.S., Foerster, E.G., Abdel-Nour, M., Philpott, D.J., and Girardin, S.E. (2019). NOD1 and NOD2 in inflammation, immunity and disease. Arch Biochem Biophys 670, 69–81. 10.1016/j.abb.2018.12.022.

65. Trindade, B.C., and Chen, G.Y. (2020). NOD1 and NOD2 in inflammatory and infectious diseases. Immunol Rev 297, 139–161. 10.1111/imr.12902.

66. Neil, J.A., Matsuzawa-Ishimoto, Y., Kernbauer-Holzl, E., Schuster, S.L., Sota, S., Venzon, M., Dallari, S., Galvao Neto, A., Hine, A., Hudesman, D., et al. (2019). IFN-I and IL-22 mediate protective effects of intestinal viral infection. Nat Microbiol 4, 1737–1749. 10.1038/s41564-019-0470-1.

67. Sugimoto, K., Ogawa, A., Mizoguchi, E., Shimomura, Y., Andoh, A., Bhan, A.K., Blumberg, R.S., Xavier, R.J., and Mizoguchi, A. (2008). IL-22 ameliorates intestinal inflammation in a mouse model of ulcerative colitis. J Clin Invest 118, 534–544. 10.1172/JCI33194.

68. Chu, H., Khosravi, A., Kusumawardhani, I.P., Kwon, A.H., Vasconcelos, A.C., Cunha, L.D., Mayer, A.E., Shen, Y., Wu, W.L., Kambal, A., et al. (2016). Gene-microbiota interactions contribute to the pathogenesis of inflammatory bowel disease. Science 352, 1116–1120. 10.1126/science.aad9948.

69. Wu, W.H., Kim, M., Chang, L.C., Assie, A., Saldana-Morales, F.B., Zegarra-Ruiz, D.F., Norwood, K., Samuel, B.S., and Diehl, G.E. (2022). Interleukin-1beta secretion induced by mucosa-associated gut commensal bacteria promotes intestinal barrier repair. Gut Microbes 14, 2014772. 10.1080/19490976.2021.2014772.

70. Seo, S.U., Kuffa, P., Kitamoto, S., Nagao-Kitamoto, H., Rousseau, J., Kim, Y.G., Nunez, G., and Kamada, N. (2015). Intestinal macrophages arising from CCR2(+) monocytes control pathogen infection by activating innate lymphoid cells. Nat Commun 6, 8010. 10.1038/ncomms9010.

71. Zhou, L., Chu, C., Teng, F., Bessman, N.J., Goc, J., Santosa, E.K., Putzel, G.G., Kabata, H., Kelsen, J.R., Baldassano, R.N., et al. (2019). Innate lymphoid cells support regulatory T cells in the intestine through interleukin-2. Nature 568, 405–409. 10.1038/s41586-019-1082-x.

72. Inohara, N., Ogura, Y., Fontalba, A., Gutierrez, O., Pons, F., Crespo, J., Fukase, K., Inamura, S., Kusumoto, S., Hashimoto, M., et al. (2003). Host recognition of bacterial muramyl dipeptide mediated through NOD2. Implications for Crohn’s disease. J Biol Chem 278, 5509–5512. 10.1074/jbc.C200673200.

73. Hugot, J.P., Chamaillard, M., Zouali, H., Lesage, S., Cezard, J.P., Belaiche, J., Almer, S., Tysk, C., O’Morain, C.A., Gassull, M., et al. (2001). Association of NOD2 leucine-rich repeat variants with susceptibility to Crohn’s disease. Nature 411, 599–603. 10.1038/35079107.

74. Murano, T., Okamoto, R., Ito, G., Nakata, T., Hibiya, S., Shimizu, H., Fujii, S., Kano, Y., Mizutani, T., Yui, S., et al. (2014). Hes1 promotes the IL-22-mediated antimicrobial response by enhancing STAT3-dependent transcription in human intestinal epithelial cells. Biochem Biophys Res Commun 443, 840–846. 10.1016/j.bbrc.2013.12.061.

75. Winter, S.E., Winter, M.G., Xavier, M.N., Thiennimitr, P., Poon, V., Keestra, A.M., Laughlin, R.C., Gomez, G., Wu, J., Lawhon, S.D., et al. (2013). Host-derived nitrate boosts growth of E. coli in the inflamed gut. Science 339, 708–711. 10.1126/science.1232467.

76. Lee, J.Y., Tsolis, R.M., and Baumler, A.J. (2022). The microbiome and gut homeostasis. Science 377, eabp9960. 10.1126/science.abp9960.

77. Shouval, D.S., Biswas, A., Kang, Y.H., Griffith, A.E., Konnikova, L., Mascanfroni, I.D., Redhu, N.S., Frei, S.M., Field, M., Doty, A.L., et al. (2016). Interleukin 1beta Mediates Intestinal Inflammation in Mice and Patients With Interleukin 10 Receptor Deficiency. Gastroenterology 151, 1100–1104. 10.1053/j.gastro.2016.08.055.

78. Kim, D., Kim, Y.G., Seo, S.U., Kim, D.J., Kamada, N., Prescott, D., Chamaillard, M., Philpott, D.J., Rosenstiel, P., Inohara, N., and Nunez, G. (2016). Nod2-mediated recognition of the microbiota is critical for mucosal adjuvant activity of cholera toxin. Nat Med 22, 524–530. 10.1038/nm.4075.

79. Martin, P.K., Marchiando, A., Xu, R., Rudensky, E., Yeung, F., Schuster, S.L., Kernbauer, E., and Cadwell, K. (2018). Autophagy proteins suppress protective type I interferon signalling in response to the murine gut microbiota. Nat Microbiol 3, 1131–1141. 10.1038/s41564-018-0229-0.

80. Matsuzawa-Ishimoto, Y., Yao, X., Koide, A., Ueberheide, B.M., Axelrad, J.E., Reis, B.S., Parsa, R., Neil, J.A., Devlin, J.C., Rudensky, E., et al. (2022). The gammadelta IEL effector API5 masks genetic susceptibility to Paneth cell death. Nature 610, 547–554. 10.1038/s41586-022-05259-y.

81. Bokulich, N.A., Kaehler, B.D., Rideout, J.R., Dillon, M., Bolyen, E., Knight, R., Huttley, G.A., and Gregory Caporaso, J. (2018). Optimizing taxonomic classification of marker-gene amplicon sequences with QIIME 2’s q2-feature-classifier plugin. Microbiome 6, 90. 10.1186/s40168-018-0470-z.

82. Lozupone, C., Lladser, M.E., Knights, D., Stombaugh, J., and Knight, R. (2011). UniFrac: an effective distance metric for microbial community comparison. ISME J 5, 169–172. 10.1038/ismej.2010.133.

83. Vazquez-Baeza, Y., Pirrung, M., Gonzalez, A., and Knight, R. (2013). EMPeror: a tool for visualizing high-throughput microbial community data. Gigascience 2, 16. 10.1186/2047-217X-2-16.

84. Dallari, S., Heaney, T., Rosas-Villegas, A., Neil, J.A., Wong, S.Y., Brown, J.J., Urbanek, K., Herrmann, C., Depledge, D.P., Dermody, T.S., and Cadwell, K. (2021). Enteric viruses evoke broad host immune responses resembling those elicited by the bacterial microbiome. Cell Host Microbe 29, 1014–1029 e1018. 10.1016/j.chom.2021.03.015.

85. Jang, K.K., Kaczmarek, M.E., Dallari, S., Chen, Y.H., Tada, T., Axelrad, J., Landau, N.R., Stapleford, K.A., and Cadwell, K. (2022). Variable susceptibility of intestinal organoid-derived monolayers to SARS-CoV-2 infection. PLoS Biol 20, e3001592. 10.1371/journal.pbio.3001592.

